# Single cell transcriptome analysis reveals an essential role for Gata2b in hematopoietic lineage decisions in zebrafish

**DOI:** 10.1101/753178

**Authors:** Emanuele Gioacchino, Cansu Koyunlar, Hans de Looper, Madelon de Jong, Tomasz Dobrzycki, Remco Hoogenboezem, Joke Peulen, Dennis Bosch, Paulette van Strien, Martin E van Royen, Pim J French, Eric Bindels, Rui Monteiro, Kirsten J Gussinklo, Ivo P Touw, Emma de Pater

**Affiliations:** Department of Hematology, Erasmus MC, Rotterdam, The Netherlands; Department of Pathology, Cancer Treatment Screening Facility, Erasmus MC Optical Imaging Centre, Erasmus MC, Rotterdam, The Netherlands; Department of Neurology, Cancer Treatment Screening Facility, Erasmus MC, Rotterdam, The Netherlands; Molecular Haematology Unit, Weatherall Institute of Molecular Medicine, John Radcliffe Hospital, University of Oxford, Oxford OX3 9DS, UK and BHF Centre of Research Excellence, Oxford, UK; Institute of Cancer and Genomic Sciences, University of Birmingham, Birmingham B152TT, UK and BHF Centre of Research Excellence, Oxford, UK

**Keywords:** Gata2b, zebrafish, single cell RNA sequencing, hematopoietic stem- and progenitor cells, myeloid lineage differentiation

## Abstract

Hematopoietic stem cells (HSCs) are tightly controlled to keep a balance between myeloid and lymphoid cell differentiation. Gata2 is a pivotal hematopoietic transcription factor required for HSC generation and maintenance. We generated a zebrafish mutant for the mammalian *Gata2* orthologue, *gata2b*. We found that in adult zebrafish, *gata2b* is required for both neutrophilic- and monocytic lineage differentiation. Single cell transcriptome analysis revealed that the myeloid defect present in Gata2b deficient zebrafish arise in the most immature hematopoietic stem and progenitor cell (HSPC) compartment and that this population is instead committed towards the lymphoid and erythroid lineage. Taken together, we find that Gata2b is vital for the fate choice between the myeloid and lymphoid lineages.

## Introduction

Hematopoietic stem cells (HSCs) have the capacity to self renew and to generate all lineages of the hematopoietic system (Sawai et al., 2016). Importantly, the HSC pool is a heterogeneous population of cells which are tightly controlled to maintain a balance between myeloid and lymphoid cell commitment (Dykstra et al., 2007; Muller-Sieburg et al., 2004; Sanjuan-Pla et al., 2013; Yamamoto et al., 2013).

Factors like aging or the microenvironment affect the lineage choice of HSCs (Gekas and Graf, 2013; Pinho et al., 2018). However, little is known about cell-intrinsic regulators of the fate choice between myeloid and lymphoid lineage differentiation. As defects in lineage differentiation contribute to a variety of hematological diseases such as bone marrow failure syndromes, myelodysplastic syndromes (MDS) and acute myeloid leukemia (AML), it is vital to understand the regulation of lineage commitment in HSPCs.

The adult hematopoietic system is generated from the definitive wave of embryonic hematopoiesis and the first HSCs transdifferentiate from specialized hemogenic endothelium (HE) of the dorsal aorta at embryonic day (E)10.5. This transdifferentiation takes place in the aorta-gonad-mesonephros region (AGM) through a process known as endothelial-to-hematopoietic transition (EHT). This process is highly conserved among species (Bertrand et al., 2010a; Boisset et al., 2010; Jaffredo et al., 1998; Kissa and Herbomel, 2010; Medvinsky and Dzierzak, 1996).

In mouse and human, Gata2 is required for embryonic HSC generation (de Pater et al., 2013; Gao et al., 2013; Huang et al., 2015; Kang et al., 2018). Germline deletion of *Gata2* in mice results in embryonic lethality at E10, just before the generation of the first HSCs (Tsai et al., 1994). Gata2 expression is tightly regulated during distinct stages of embryonic development and Gata2 plays crucial roles in the specification of hemogenic endothelium and the generation and maintenance of HSCs (de Pater et al., 2013; Gao et al., 2013; Johnson et al., 2015; Mehta et al., 2017; Snow et al., 2010). Gata2 has also been implicated in myeloid differentiation, supported by studies showing that hematopoietic stem and progenitor cells (HSPCs) from mice with heterozygous loss-of-function mutations in Gata2 have reduced granulocytic and monocytic potential (Ling et al., 2004; Rodrigues et al., 2008; Rodrigues et al., 2005) and Gata2 is expressed in several myeloid cell populations (Kauts et al., 2018; Zon et al., 1993). A role for Gata2 in myeloid/lymphoid lineage differentiation is further supported by that the fact that overexpression of Gata2 resulted in a block in lymphoid differentiation (Nandakumar et al., 2015) and that in human embryonic stem cell cultures GATA2 knockout cells are incapable of generating granulocytes (Huang et al., 2015; Kang et al., 2018).

Due to its embryonic lethality in mice, *in vivo* studies of the function of Gata2 in lineage commitment and differentiation are cumbersome. Also, the various roles Gata2 plays during embryonic development make it difficult to dissect its exact function during the processes of HE specification, EHT and lineage differentiation. Zebrafish are an ideal *in vivo* model to study the function of Gata2 in hematopoiesis, because the embryonic stages do not depend on the hematopoietic system for survival (Driever et al., 1996; Haffter et al., 1996). Moreover, zebrafish embryonic hematopoietic development resembles that of mammals. For instance, like in mice, the first HSCs are generated in the AGM region and then amplified in the caudal hematopoietic region (CHT), which functions as the fetal liver homologue in zebrafish (Bertrand et al., 2010a; Bertrand et al., 2010b; Ciau-Uitz et al., 2014; Kissa and Herbomel, 2010; Tamplin et al., 2015; Warga et al., 2009). In zebrafish, adult hematopoiesis takes place in the kidney marrow instead of the bone marrow (Traver et al., 2003). The kidney marrow contains all hematopoietic lineages and morphology of hematopoietic cells is comparable to human hematopoietic cells (Davidson and Zon, 2004).

Zebrafish have two orthologues of Gata2; Gata2a and Gata2b. Previous studies have shown that *gata2b* is more prominently expressed in HSPCs whereas Gata2a is mainly expressed in the vasculature. knockdown of *gata2b* severely reduces definitive hematopoiesis during embryonic stages and lineage analysis revealed that all definitive hematopoietic cells once expressed *gata2b* (Butko et al., 2015), indicating that Gata2b is the predominant Gata2 orthologue required for the maintenance of hematopoietic stem cells.

The present study features the role of Gata2b in hematopoietic cell development. We show that HSPC generation is not affected in *gata2b^-/-^* embryos, but that the number of HSPCs is reduced at 76 hours post fertilization (hpf) in the CHT after the first amplification phase. In addition, we show that in adult zebrafish, Gata2b is required for differentiation of the neutrophil and monocyte lineage. Prospective isolation and transcriptome analysis of the phenotypic HSC compartment showed overexpression of several lymphoid lineage markers in Gata2b deficient cells. Furthermore, single cell transcriptome analysis revealed that the majority of Gata2b deficient immature HSPCs have an increased proliferation signature and a lymphoid lineage transcriptional profile. Together these data show a pivotal role for Gata2b in HSPC expansion during embryogenesis and in adults in the lineage choice between myeloid and lymphoid lineage differentiation.

## Results

### Gata2b is dispensible for generation of hematopoietic stem cells

To generate Gata2b mutants, we used CRISPR/Cas9 to target the third exon of *gata2b* located upstream of the DNA and protein binding zinc fingers (znf) which are encoded by the fifth and the sixth exon (Figure 1A). A 28 bp integration was introduced in the third exon leading to a frameshift truncation from amino acid 185 (Figure 1B, C, D). Hereafter, we refer to this mutant as *gata2b^-/-^*.

**Figure 1.**
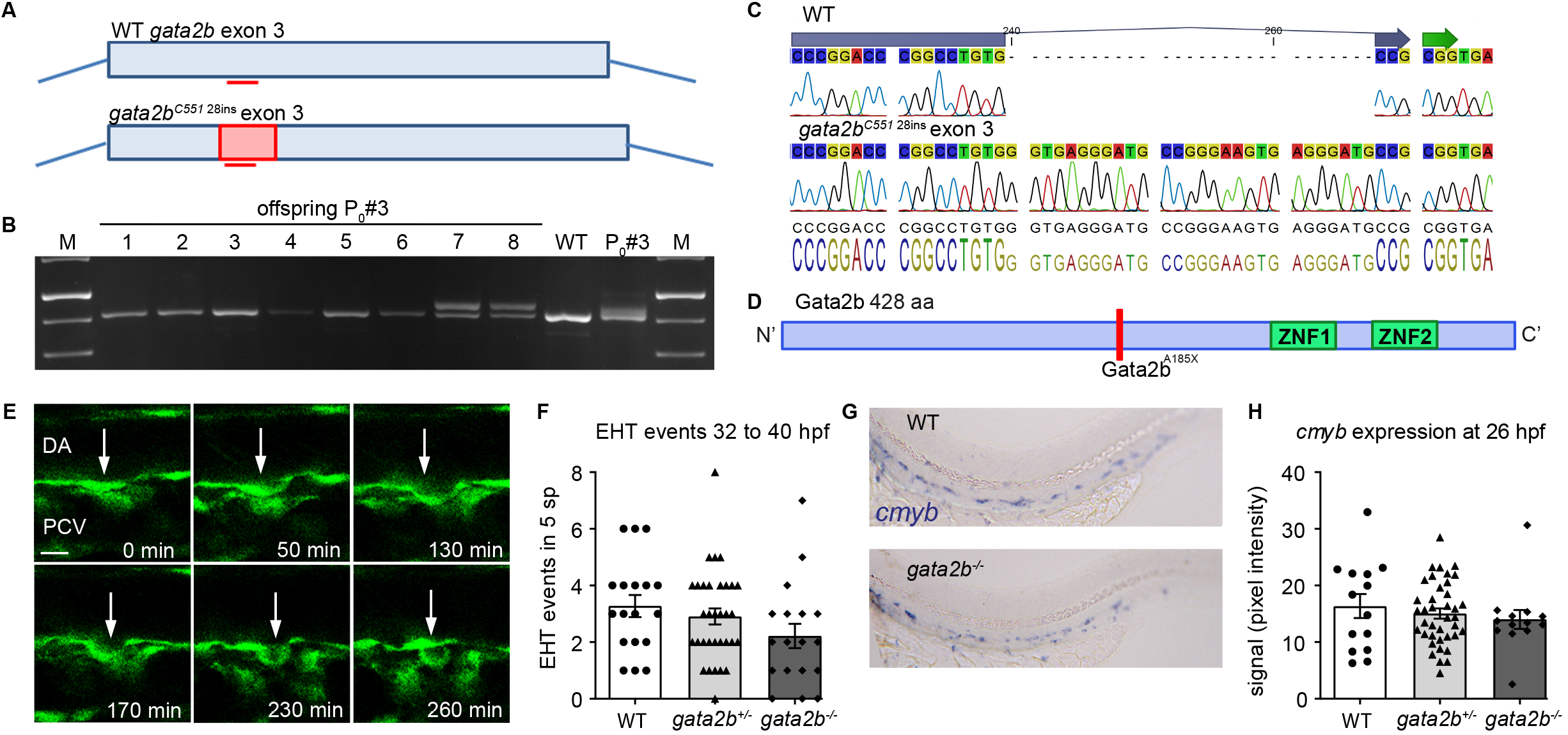
Newly generated Gata2b mutant zebrafish does not show defects in HSPC generation. A) Schematic representation of the CRISPR strategy targeting exon 3 of *gata2b* and the 28 nt integration in *gata2b* mutants. B) Gel picture showing genotyping PCR of founder 3 and the F1 with a 28 bp integration in embryo 7 and 8. C) Alignment of sequencing data of WT *gata2b* exon 3 where the location of the guide is indicated in the blue arrow on top of the sequence and sequencing data from *gata2b^-/-^* DNA showing a 28 nucleotide integration. D) Gata2b mutation leading to a STOP codon abrogating the protein before the DNA and protein binding znk fingers. E) Example of EHT event from WT *Tg(fli1a:eGFP)* transgenic zebrafish. Time indicated at the bottom right corner in minutes. Scalebar represents 10 μm. arrow indicates endothelial cell undergoing hematopoietic transition. F) Quantitation of EHT events between 32-40 hpf in WT, *gata2b^+/-^* and *gata2b^-/-^* embryos in an area of 5 sp in the AGM region. Each dot represents one embryo. G) Representative image of *cmyb* expression in WT and *gata2b^-/-^* embryos at 26 hpf. H) Quantitation of *cmyb* signal intensity relative to background in WT, *gata2b^+/-^* and *gata2b^-/-^* embryos in the AGM region. P_0_ = founder, Bp = basepair, aa = amino acid, EHT = endothelial to hematopoietic transition, DA = dorsal aorta, PVC = posterior cardinal vein, sp = somite pair, hpf = hours port fertilization. Error bars represent SEM.

Because HSCs are generated through EHT, we first analysed the number of EHT events in *gata2b^-/-^* embryos. We visualized EHT using *Tg(Fli1a:GFP)*, marking all endothelial cells including hemogenic endothelium, and quantified the cells that underwent EHT in a 5 somite pair (sp) region using time-lapse confocal microscopy (Figure 1E, F). *gata2b^-/-^* embryos showed a trend towards reduced EHT events between 32-40 hours post fertilization (hpf) (P = 0.077, Figure 1F and Table 1) but no significant difference was detected in EHT events between WT and Gata2b deficient animals. HE is marked by *cmyb* at 26 hours post fertilization (hpf) (Bertrand et al., 2010a; Bertrand et al., 2008). To enumerate HE we quantified *cmyb* expression by measuring pixel intensity of the *in situ* hybridization (*ish*) staining compared to background (Dobrzycki et al., 2018) in the AGM region at 26 hpf, the earliest time of EHT events in zebrafish (Figure 1G, H). Expression of *cmyb* is not altered between WT and Gata2b deficient
embryos at 26 hpf, indicating that specification of hemogenic endothelium occurs normally in the absence of Gata2b (Figure 1H and Table 1). Taken together, these experiments indicate that quantitatively HSPC generation through EHT is not or only marginally affected in *gata2b^-/-^* embryos.

**Table 1.** Embryonic hematopoietic cell quantitations

| Area and time point | WT | *gata2b*<sup>+/-</sup> | *gata2b*<sup>-/-</sup> |
| --- | --- | --- | --- |
| **EHT events** |  |  |  |
| AGM (32 to 40hpf) | 3.3 ± 0.4 (n=18) | 2.9 ± 0.3 (n=33) | 2.2 ± 0.4 (n=18) |
| **CD41:GFP<sup>+</sup>flt:RFP<sup>+</sup> cells** |  |  |  |
| CHT (52-54hpf) | 8.5 ± 1.7 (n=8) | 9.0 ± 1.5 (n=20) | 11.9 ± 1.0 (n=7) |
| CHT (58-60hpf) | 25.2 ± 3.2 (n=16) | 28.7 ± 1.9 (n=28) | 23.3 ± 2.9 (n=12) |
| CHT (76-58hpf) | 55.5 ± 3.8 (n=13) | 43.4 ± 5.6 (n=12) | 36.5 ± 3.0 (n=14)** |
| ***cmyb* expression intensity** |  |  |  |
| AGM (24 hpf) | 33.0 ± 6.3 (n=14) | 28.5 ± 4.5 (n=39) | 30.6 ± 2.6 (n=13) |
| AGM (33 hpf) | 32.9 ± 10.9 (n=16) | 30.9 ± 7.9 (n=31) | 28.5 ± 6.5 (n=28)** |
| CHT (56 hpf) | 10.5 ± 2.5 (n=19) | 12.8 ± 0.5 (n=41) | 4.2 ± 1.1 (n=14)** |
| CHT (76 hpf) | 21.3 ± 7.4 (n=19) | 20.8 ± 1.1 (n=43)** | 16.1 ± 2.0 (n=15)* |
| ***Tg(mpeg:GFP)*<sup>+</sup> cells** |  |  |  |
| CHT (54hpf) | 513.9 ± 23.7 (n=18) | 564.9 ± 23.5 (n=26) | 545.9 ± 33.3 (n=14) |
| ***Tg(mpx:GFP)*<sup>+</sup> cells** |  |  |  |
| CHT (75hpf) | 227.5 ± 14.4 (n=11) | 221.1 ± 8.6 (n=26) | 235.0 ± 15.5 (n=6) |
| ***Tg(LCK:GFP)*<sup>+</sup> pixel number** |  |  |  |
| Thymus (5dpf) | 2093 ± 186.2 (n=9) | 2449 ± 165.5 (n=18) | 2354 ± 206.8 (n=17) |

The data are mean ± SEM. n= number of zebrafish embryos used in analysis. AGM; aorta-gonad-mesonephros region, CHT; caudal hematopoietic tissue. \*P< 0.05, \*\*P< 0.01. If data are normally distributed we used One-way ANOVA with Tukey post test. If data are not normally distributed we used Kruskal-Wallis with Dunn's post test.

### Gata2b is required for embryonic definitive hematopoiesis

To analyse the effect of Gata2b deficiency on definitive hematopoiesis, we used a combination of phenotypic markers to investigate transcriptional integrity of HSPCs. HSPCs, which are generated in the AGM region, are marked by *cmyb* expression and can be visualized by *in situ* hybridization (North et al., 2007). Although the generation of HSPCs is not affected by Gata2b deletion (Figure 1H), from 33 hpf onward the expression of *cmyb* is significantly reduced in the AGM region of *gata2b^-/-^* embryos compared to WT (Figure 2A,B and Table 1). Similarly, *cmyb* expression is reduced in the CHT of *gata2b^-/-^* embryos at 56 hpf, when HSPCs are expanded (Figure 2C,D and Table 1). From 44 hpf onward, HSPCs are marked by the coexpression of *Tg(CD41:GFP)* and the arterial marker *Tg(Flt:RFP)* as definitive HSPCs are derived from arteries (Bussmann et al., 2010; Ma et al., 2011) (Figure 2E). CD41:GFP^+^Flt:RFP^+^ cells were unchanged at 52-54 hpf in *gata2b^-/-^* embryos compared to WT (Figure 2F and Table 1) and 58-60 hpf in *gata2b^-/-^* embryos compared to WT (Figure 2G and Table 1). However, at 76 hpf, CD41:GFP^+^Flt:RFP^+^ cells were reduced in the CHT in *gata2b^-/-^* embryos compared to WT (Figure 2H and Table 1). From 52 hpf to 76 hpf, the number of HSPCs increased rapidly in the CHT in WT embryos (6.5 fold) whereas a smaller increase in *gata2b^-/-^* embryos was observed (3.1 fold from 52 hpf to 76 hpf). The discordance between the CD41 marker analysis and the *cmyb* expression indicates that the lower expression of *cmyb* in *gata2b^-/-^* HSPCs anticipates the decrease in phenotypic HSPCs. These findings indicate that during the amplification phase of *gata2b^-/-^* embryos there might be either a selection of a subclass of definitive hematopoietic cells or a general impairment in cell division.

**Figure 2.**
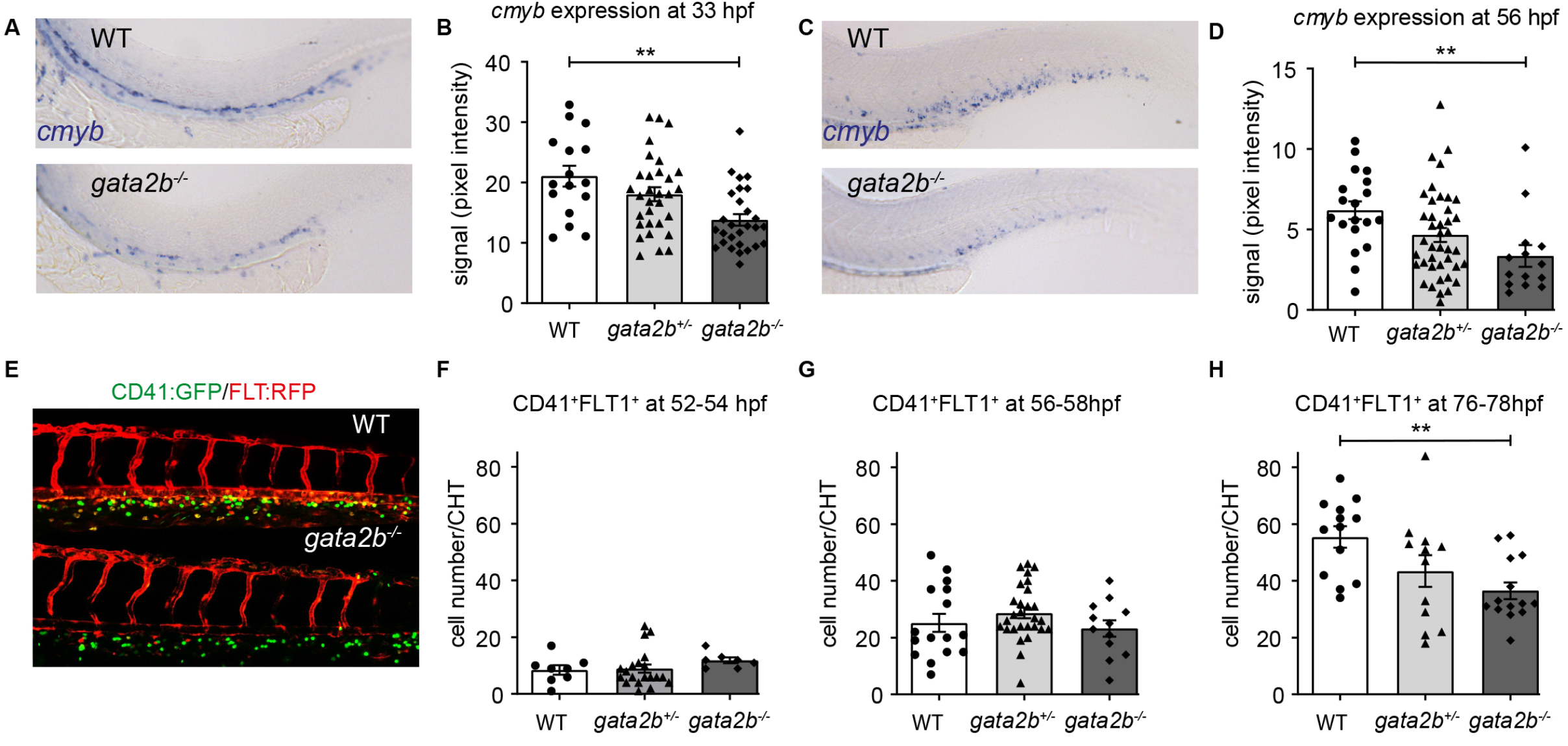
*Gata2b^-/-^* embryos show a reduction in hematopoietic stem and progenitor marker expression. (A) Representative example of *cmyb* expression in WT and *gata2b^-/-^* embryos at 33 hpf in the AGM region and C) 56 hpf in the CHT B) Quantitation of signal intensity relative to background cells in WT, *gata2b^+/-^* and *gata2b^-/-^* embryos at 33 hpf where each dot represents one embryo and D) in 56 hpf in the CHT. E) example of *Tg(CD41:GFP); Tg(Flt:RFP)* expression in the CHT of WT and *gata2b^-/-^* embryos at 76 hpf. F-G) quantitation of GFP^+^RFP^+^ cells in WT, *gata2b^+/-^* and *gata2b^-/-^* embryos where each dot represents one embryo at F) 52-54 hpf, G) 56-58 hpf, and H) 76-78 hpf. hpf = hours post fertilization, CHT = caudal hematopoietic tissue. * = P < 0.05, ** = P < 0.01. Error bars represent SEM.

### Lack of Gata2b leads to reduced myeloid differentiation, lymphoid bias and an accumulation of proerythroblasts in adult kidney marrow

Maternal-zygotic *gata2b^-/-^* embryos survive to adulthood in mendelian ratios (Figure S2A, B). Because Gata2 has been implicated in myeloid differentiation in the mouse (Rodrigues et al., 2008), we asked whether hematopoietic differentiation was affected during embryonic development. Lymphoid and myeloid differentiation was normal in Gata2b deficient embryos as determined by lineage marker analysis (Figure S1). Next, we investigated hematopoietic lineage differentiation in the adult *gata2b^-/-^* kidney marrow (KM) by scatter profile analysis, transgenic marker analysis and morphological analysis (Ellett et al., 2011; Ma et al., 2011; Renshaw et al., 2006; Traver et al., 2003).

While *gata2b^-/-^* embryos did not show signs of altered lineage differentiation (Figure S1), scatter profile analysis of adult *gata2b^-/-^* zebrafish KM showed a significant reduction in the myeloid gate of 3-5 month post fertilization (mpf) zebrafish (Figure 3A, B and Table S1) and a relative increase in the HSPC and lymphocytes gate (31.9% ± 1.6 vs 19.4% ± 1.2, *gata2b^-/-^* vs WT)(Figure 3A, B and Table S1).

**Figure 3.**
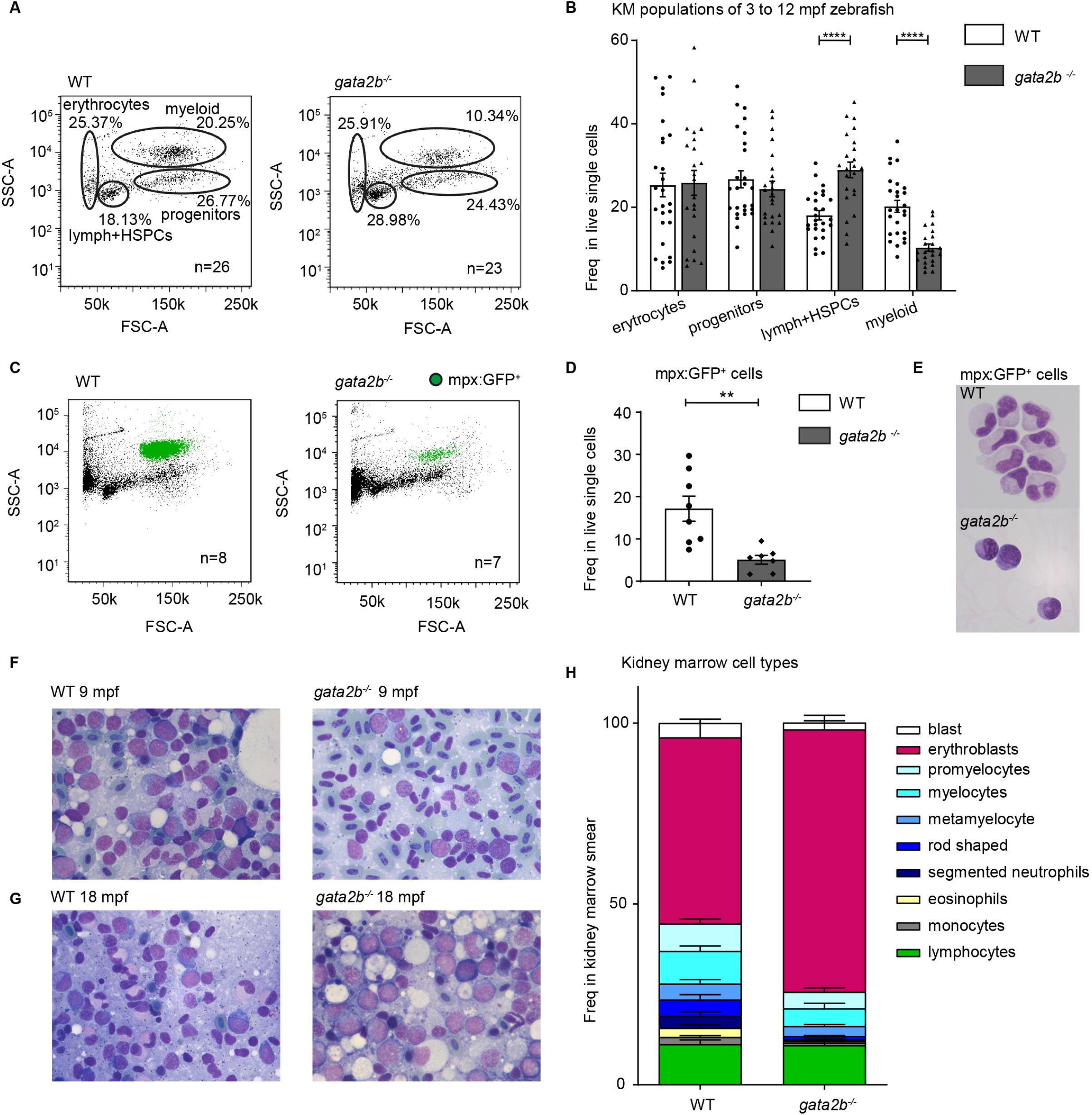
Gata2b deficiency results in decreased myeloid differentiation in adult zebrafish KM. A) Gating strategy of FACS analysis of whole kidney marrow of WT and *gata2b^-/-^* zebrafish. B) Quantitation of erythroid, myeloid, lymphoid and HSPC and progenitor gates as percentages of single viable cells of KM harvested from 3-12 mpf zebrafish. Each dot represents kidney marrow analysis of one zebrafish. C) FSC-A/SSC-A representation of whole kidney marrow in *Tg(mpx:GFP)* WT and *gata2b^-/-^* zebrafish with the GFP positive population indicated in green. D) quantitation of GFP^+^ percentage in single viable cells. Each dot represents kidney marrow analysis of one zebrafish. E) Cytospin, stained with MGG, of *Tg(mpx:GFP)^+^* sorted cells of WT and *gata2b^-/-^* zebrafish F) kidney marrow smear of WT and *gata2b^-/-^* zebrafish at 9 mpf and G) 18 mpf. H) Quantitation of the frequencies of cell types in kidney marrow smears after morphological differentiation taken form 9 - 18 mpf zebrafish. * = P < 0.05, ** = P < 0.01, *** = P < 0.001, **** = P < 0.0001. KM = kidney marrow, mpf = months post fertilization, SSC = side scatter, FSC = forward scatter, lymph = lymphocytes, Error bars represent SEM.

To specifically address which lineage was affected by the loss of Gata2b, *gata2b^-/-^* zebrafish were crossed with several transgenic lines, marking specific hematopoietic cell populations. Analysis of *Tg(mpx:GFP)* zebrafish showed a severe reduction in mature neutrophils in *gata2b^-/-^* kidney marrow (Figure 3C, D and Table S1). Sorted mpx:GFP+ cells from *Tg(mpx:GFP)* WT and *gata2b^-/-^* zebrafish showed that the remaining *gata2b^-/-^* mpx:GFP+ cells show a more immature neutrophil morphology (Figure 3E), indicating that Gata2b is required for normal neutrophil differentiation. Additionally, morphological analysis of kidney marrow smears taken from 9 month old (Figure 3F) and 18 month old (Figure 3G) zebrafish kidneys showed that *gata2b^-/-^* zebrafish developed an erythroid lineage bias, ultimately leading to an accumulation of pro-erythroblasts at 18 months post fertilization as detected by the quantitation of kidney marrow smears (Figure 3H). Furthermore, the kidney marrow smears showed an almost complete absence of monocytes, segmented neutrophils and the more immature rod shaped neutrophils, confirming that myeloid differentiation was severely compromised (Figure 3H).

### Adult gata2b^-/-^ phenotypic HSPCs show upregulation of lymphoid transcription factors

Because Gata2 is also required for HSC maintenance in mice (de Pater et al., 2013; Ling et al., 2004; Rodrigues et al., 2005), we hypothesized that the lineage differentiation defect observed in *gata2b^-/-^* zebrafish arises in the stem cell compartment. In adult zebrafish, the HSC population is most stringently marked by CD41:GFP^low^ expression, excluding most progenitors (Ma et al., 2011). To investigate whether the *gata2b^-/-^* HSPC population shows lineage bias, CD41:GFP^low^ cells were prospectively isolated from WT and *gata2b^-/-^* zebrafish kidney marrow followed by transcriptome analysis (Figure 4A, B). WT and Gata2b deficient zebrafish had comparable numbers of CD41:GFP^low^ cells (Figure 4C) but analysis of the transcriptome of *gata2b^-/-^* CD41:GFP^low^ cells by RNA sequencing showed 6 gene sets that were significantly downregulated in *gata2b^-/-^* CD41:GFP^low^ cells. Some of these gene sets are related to chromatin regulation (Figure S3A-F). In-depth analysis of the differentially regulated genes in *gata2b^-/-^* CD41:GFP^low^ cells revealed that the lymphoid transcription factor *Ikaros2 (ikzf2)* and the transcriptional co-activator *B-cell lymphoma 3 protein (Bcl3)* are significantly upregulated in *gata2b^-/-^* CD41:GFP^low^ expressing cells (Figure 4D). The Ikaros family of transcription factors in mice activate lymphoid differentiation (Kim et al., 1999) and in mice, *Ikzf2* expression is associated with a block in myeloid lineage differentiation (Park et al., 2019). *Bcl3* regulates transcription by associating with NFκB and is important for B-cell proliferative responses and involved in T-helper cell differentiation (reviewed in (Herrington and Nibbs, 2016). The upregulation of these lymphoid program genes in *gata2b^-/-^* CD41:GFP^low^ cells together with the forward and side scatter differences suggests a lymphoid bias in *gata2b^-/-^* HSPCs.

**Figure 4.**
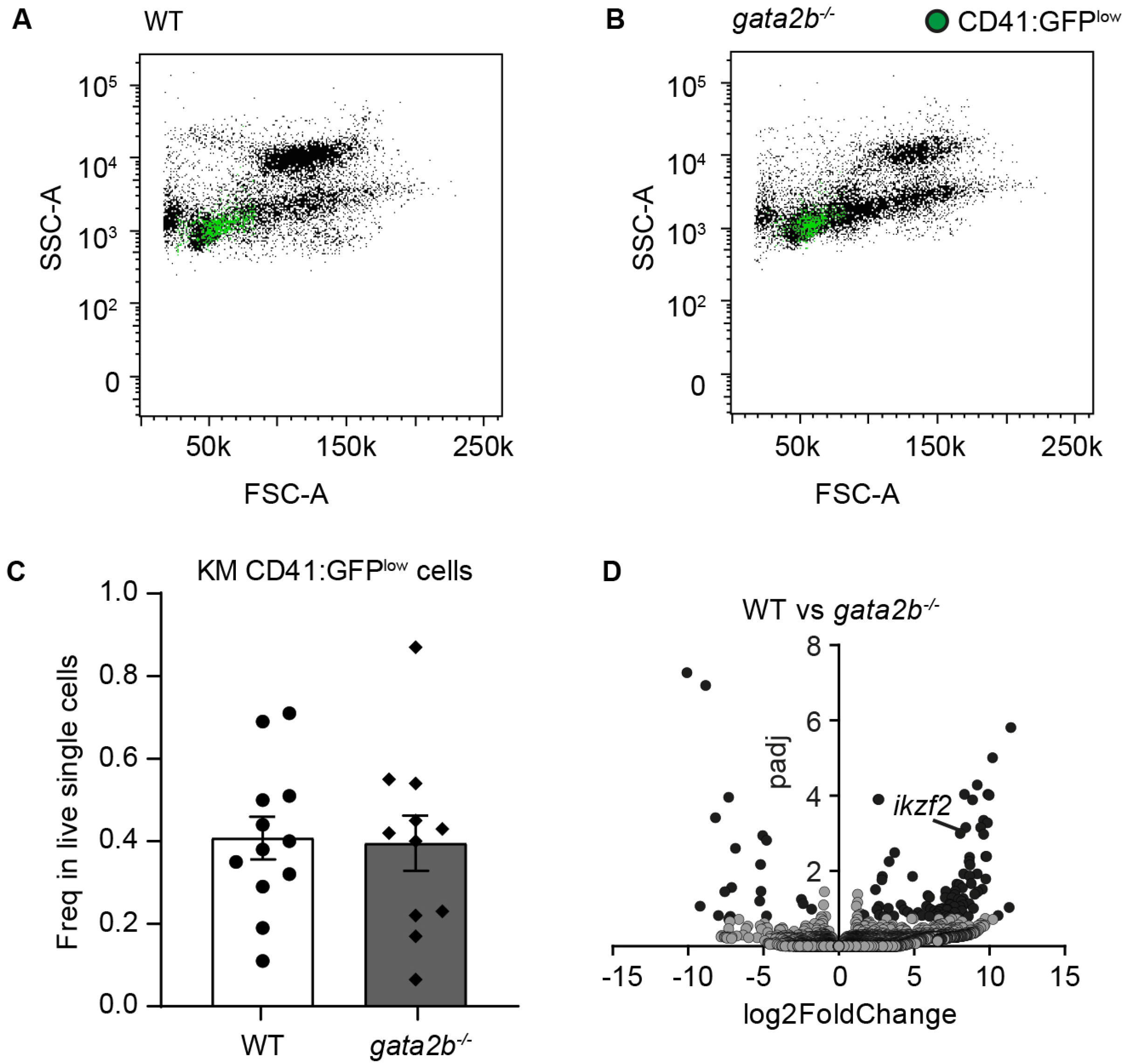
*gata2b^-/-^* phenotypic HSPCs show an increase in *ikzf2* expression. A) FSC-A/SSC-A representation of whole kidney marrow in WT and B) *gata2b^-/-^* zebrafish with the CD41:GFP^low^ population indicated in green. C) Frequency of CD41:GFP^low^ cells in WT and *gata2b^-/-^* kidney marrow. Each dot represents KM analysis of one zebrafish. Error bars represent SEM. D) Volcano plot of differential gene expression of *gata2b^-/-^* vs WT CD41-GFP^low^ cells with an indication of the differential gene expression of *ikzf2*. Genes with a significant (P<0.01) differential gene expression (log2Fold change > 1) are depicted in black. SSC = side scatter, FSC = forward scatter, KM = kidney marrow.

### Single cell RNAseq shows a lymphoid bias at the expense of the neutrophil/monocytic lineage in gata2b^-/-^ kidney marrow

The scatter profile analysis showed that the lymphoid population was greatly increased at the expense of the myeloid population in *gata2b^-/-^* zebrafish relative to wild type controls (Figure 3A). To confirm the lineage bias in *gata2b^-/-^* kidney marrow and to clarify the underlying transcriptional profile we performed single cell transcriptional profiling of all hematopoietic progenitors in zebrafish KM. In this analysis we addressed the following questions:

1. Is the lineage differentiation occurring at the level of HSPCs, more commited progenitors or is it a defect in differentiated cells? 2) Are *gata2b^-/-^* kidney marrow cells merely lineage biased or do they acquire an aberrant transcriptome?

Because, specific progenitor populations cannot be distinguished in zebrafish using cell surface markers, we sorted the entire progenitor population including the lymphoid cells of *Tg(CD41:GFP)* WT and *gata2b^-/-^* zebrafish (Figure S4A-C). Terminally differentiated erythrocytes (nucleated in zebrafish) were excluded because their high number would mask differences in other cell types and because they turned out to be present in all gates during sorting. Also, the differentiated myeloid population was excluded. We identified 20 different cell clusters using the nearest neighbor algorithm in the R Seurat package (Butler et al., 2018). Most of the clusters express specific markers which enabled us to characterize the differentiated cell types or the lineage commitment direction. Cluster identification is described in the supplementary material (Figure 5A, B, Figure S4D, E and STAR methods).

**Figure 5.**
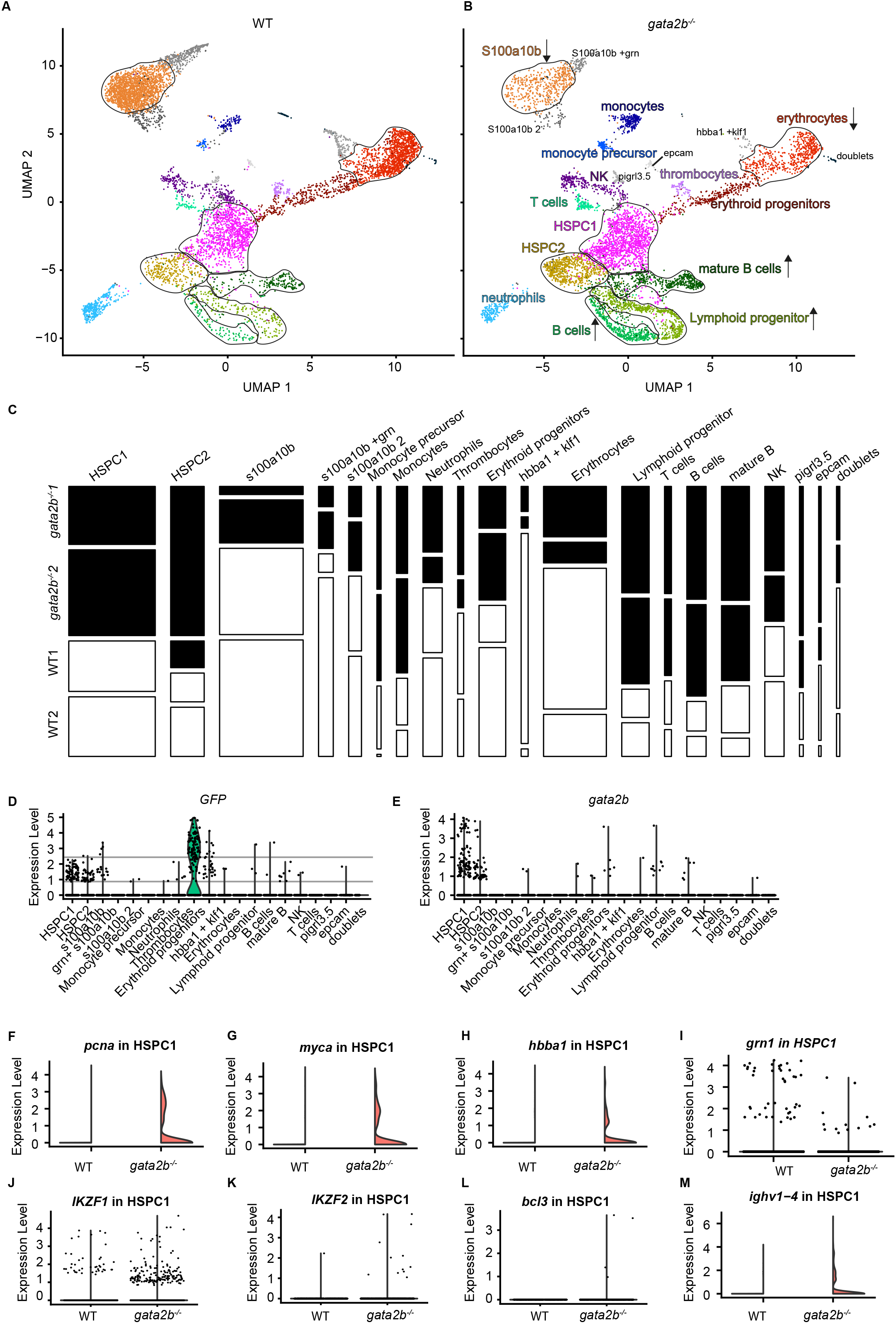
*gata2b^-/-^* cells are overrepresented in lymphoid lineage clusters and reduced in erythroid and myeloid lineage clusters compared to WT. A) Split UMAP of WT and B) *gata2b^-/-^* cells with cluster indication of enriched (arrow up) or reduced (arrow down) cell clusters in B. C) Genotype distribution of each of the clusters individually indicating the different replicate experiments. Area of the bars indicate the cell numbers in each cluster, White = WT, Black = *gata2b^-/-^*. D) Violin plot of cells with GFP gene expression from *Tg(CD41:GFP)* within the different clusters. Each dot represents one cell. The grey lines indicate GFP^low^ expression, indicating that the HSPC1 cluster contains GFP^low^ expressing cells and the thrombocyte cluster mainly contains GFP^high^ expressing cells. E) Violin plot of *gata2b* expressing cells within the different clusters. Each dot represents one cell. Most *gata2b* expressing cells are present in the HSPC1 cluster. F-M) Split violin plots of marker expression within the HSPC1 population between WT on the left and *gata2b^-/-^* cells on the right (pink) with F) *proliferating cell nuclear antigen (pcna)*, G) *MYC proto-oncogene a (myca)*, H) *hemoglobin a1 (hbba1)*, I) *granulin 1 (grn1)*, J) *ikaros family zinc finger 1 (IKZF1)*, K) *ikaros family zinc finger 2 (IKZF2), L) B cell lymphoma 3 (bcl3)* and M) *Immunoglobulin heavy variable 1-4 (ighv1-4)* expression. UMAP = Uniform manifold approximation and Projection, HSPC = Hematopoietic stem and progenitor cell.

To investigate lineage contribution between WT and *gata2b^-/-^* cells within the clusters, the proportion of these cells was analysed (Figure 5C). Five clusters were underrepresented in *gata2b^-/-^* kidney marrow, two of the erythroid lineage and three of the myeloid lineage (Figure 5A, B and C). The most striking difference was found in a large myeloid population subdivided in three clusters expressing high levels of *s100a10b* (Figure 5A-C). In human, S100A10 is highly expressed by macrophages and required for their migration (Laumonnier et al., 2006; O’Connell et al., 2010). *In situ* hybridization analysis on zebrafish sections showed that *s100a10b* is expressed in macrophages and their progenitors indicating that these cell types are most affected by Gata2b deficiency (Figure S4F-H). Additionally, *gata2b^-/-^* cells were overrepresented in the lymphoid progenitor cluster, T-cells and both B-cell clusters (Figure 5C). Therefore, single cell transcriptome analysis confirms that lineage differentiation is biased towards the lymphoid lineage at the expense of the myeloid in Gata2b deficient kidney marrow.

### Differential gene expression reveals increased proliferation and lymphoid marker expression in the gata2b^-/-^ HSPCs

The bulk transcriptome analysis on HSPCs supported the idea that the lineage skewing might occur in a progenitor or more immature population as opposed to more committed progenitors or terminally differentiated cells. We identified two HSPC cell clusters called HSPC1 and 2 (Fig. 5A and B). These clusters are marked by HSC genes like *fli1a* and *meis1b* (Athanasiadis et al., 2017; Macaulay et al., 2016; Tang et al., 2017) (Figure S4I, J), by proliferation markers (differentiated cells have low proliferation signatures) like *pcna, myca* and *mki67* (Figure S4E) and GFP from the CD41:GFP transgene (Figure 5D). Also, most *gata2b* expressing cells are found in HSPC1 and HSPC2 (Figure 5E) indicating that these populations would be most affected by loss of Gata2b. Compared to WT HSPC1s, mutant cells express higher levels of proliferation marker genes like *pcna* (Figure 5F), *myca* (Figure 5G), *hemoglobin (hbba1)* (Figure 5H), and lymphoid marker genes like *ikzf1, ikzf2, bcl3* and *ighv1-4* (Figure 5J-M); indicating that this mutant population is re-directed towards a lymphoid and erythroid transcriptional program. The *gata2b^-/-^* HSPC1 cluster contains both fewer cells expressing myeloid marker genes and an average lower expression level of the myeloid marker genes *cebpa* and *grn1* (Figure 5I and data not shown). Based on this data we conclude that the lymphoid bias in *gata2b^-/-^* zebrafish kidney marrow initiates in the most immature HSPC1.

The HSPC2 population has a similar, but more exaggerated, transcriptional profile compared to HSPC1 (Figure S5) indicating this has a more mature progenitor population compared to HSPC1. Like HSPC1, differential gene expression analysis in HSPC2 revealed a similar lymphoid bias in *gata2b^-/-^* cells (Figure S5I-M) at the expense of the myeloid program (Figure S5N-R).

### Cell fate of gata2b^-/-^ kidney marrow HSPCs is redirected to the lymphoid lineage

In *gata2b^-/-^* kidney marrow, lymphocyte populations accumulated while monocyte and granulocyte differentiation was reduced, but the cells did not show any morphological defects. Therefore, we propose that Gata2b deficiency leads to a switch in lineage choice instead of a general aberrant transcriptome in HSPCs. To further investigate the nature of the transcriptional changes in *gata2b^-/-^* HSPC1s we performed reclustering of the HSPC1 population (Figure6A, B). In this way, clustering is only based on genes that are (differentially) expressed in the HSPC1 population. Proportion analysis showed that subcluster 1 was mainly composed of WT cells and subclusters 5, 6 and 7 were mainly composed of *gata2b^-/-^* cells (Figure 6C), indicating that the differences between clusters were defined by the differences between WT and *gata2b^-/-^* cells rather then a unique myeloid or lymphoid progenitor profile. As expected, most *gata2b* expressing cells in WT are located in subcluster 1, indicating that this subcluster is most affected by Gata2b deficiency (Figure 6D). This was further strengthened by the finding that reclustering of HSPC1 did not generate subpopulations expressing only stem cell markers (Figure 6F), myeloid/erythroid markers (Figure 6E, G), proliferation markers (Figure 6H, I), or lymphoid marker expression (Figure 6J-M).

**Figure 6.**
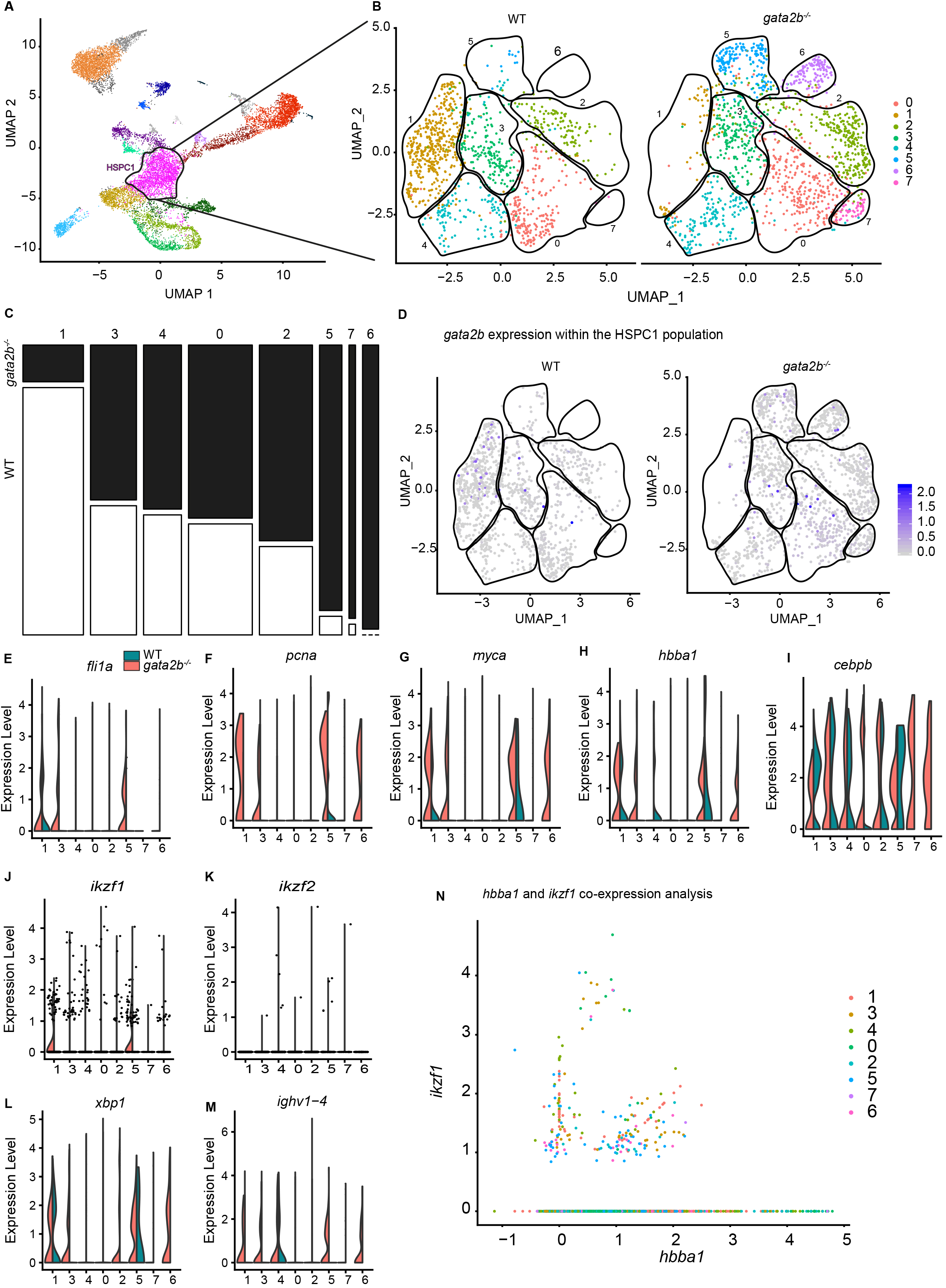
HSPC1 reclustering shows more *gata2b^-/-^* cells with erythroid and lymphoid marker expression then WT cells. A) Cluster selection for reclustering B) Reclustering of the HSPC1 population split between WT and *gata2b^-/-^* cells. C) Genotype distribution of WT and *gata2b^-/-^* HSPC1 in each of the clusters with WT cells in white and *gata2b^-/-^* cells in black. D) Feature analysis of *gata2b* expression in blue within WT on the left and gata2b^-/-^HSPC1 cells on the right. E-M)Violin plots of marker expression in the different clusters split by genotype with expression in WT cells in green and expression in *gata2b^-/-^* cells in pink. E) HSPC marker *fli1a*, proliferation markers F) *pcna* G) *myca* and lymphoid markers H) myeloid marker *cebpb* expression, I) erythroid marker *hbba1* expression, lymphoid marker expression J) *ikzf1*, K) *ikzf2*, L) *x-box protein coding 1 (xbp1)*, M) *ighv1-4*. N) Co-expression analysis of *hbba1* and *ikzf1*. Colors indicate the different subclusters of the HSPC1 population. UMAP = Uniform manifold approximation and Projection, HSPC = Hematopoietic Stem or Progenitor Cell.

Differential gene expression analysis showed that the *gata2b^-/-^* cells in cluster 1 had a lower expression of *cebpb*, a myeloid lineage marker, than the WT cells in this cluster (Figure 6E). Additionally, we found high expression of lymphoid lineage genes expressed in *gata2b^-/-^* cells (Figure 6J-M), but the fact that we also found expression of these lymphoid lineage marker genes (including variable chain immunoglobulins) in WT cells indicates that this transcriptional profile is normal (Figure 6M, subcluster 4). Moreover, in case of an aberrant transcriptome caused by the *gata2b* mutation we would expect coexpression of marker genes of multiple lineages in the same cells. Instead cells expressed either the erythroid marker *hbba1* or the lymphoid marker *ikzf1* in the HSPC1 cluster (Figure 6N). Taken together, these results led us to conclude that Gata2b deficient zebrafish HSC cell fate is redirected leading to an accumulation of lymphoid biased HSPCs and a reduction in myeloid biased HSPCs causing the lineage bias in adult zebrafish.

## Discussion

In this study, we showed that Gata2b is not vital for embryonic generation of HSPCs, but that definitive hematopoiesis is affected from 76 hpf in the caudal hematopoietic tissue. In addition, adult Gata2b deficient kidney marrow showed a lymphoid bias at the expense of the myeloid lineage based on scatter profiles and transgenic analyses. Unbiased single cell transcriptome analysis confirmed that the loss of Gata2b results in decreased myeloid differentiation and was biased towards lymphoid differentiation from 5 months post fertilization onwards. Finally, the HSPC1 compartment was the origin of the lymphoid lineage bias in *gata2b^-/-^* kidney marrow cells, due to an upregulation of the lymphoid lineage gene expression program. These data establish that Gata2b is vital for maintaining the myeloid differentiation program, and that in the absence of Gata2b, HSPCs initiate the lymphoid differentiation program instead.

The molecular mechanism controlling lineage commitment has long been thought to be regulated by stochastic variation in transcription factors (Graf and Enver, 2009). However, later reports suggested that some transcription factors have a reinforcing activity for terminal differentiation instead of lineage choice and propose that microenvironmental or upstream regulators are decisive for lineage commitment (Hoppe et al., 2016). Our results are consistent with Gata2b being an essential regulator of lineage decision in HSPCS. Both mutant and wild type HSPC1s express divergent lineage markers in the same cell. This divergent expression has been noticed before in single cell transcriptome analyses of mouse and human HSPCs (Drissen et al., 2016; Velten et al., 2017). In fact, this conflicting expression of markers might regulate lineage priming. Interestingly, Gata2 has been described to directly interact with Ikaros to regulate transcriptiption (Bottardi et al., 2013). Whether this direct interaction regulates myeloid and lymphoid bias directly or whether Gata2 acts as a repressor or a co-activator in this complex with Ikaros is as yet unclear.

Not all myeloid lineage differentiation was abrogated and few intact neutrophils remained present in Gata2b deficient HSPCs. Also, the monocyte precursor and monocyte cluster, marked by *mpeg1.1* were present and even enriched in the Gata2b deficient cells as analysed by single cell analysis. This could be explained by tissue resident macrophages in the kidney. Tissue resident macrophages are derived from embryonic erythroid-myeloid-progenitors (Bertrand et al., 2005; Herbomel et al., 1999) and myeloid embryonic differentiation was not affect in *gata2b^-/-^* zebrafish. Kindey marrow cells from 18 month old *gata2b^-/-^* zebrafish show an accumulation of pro-erythroblasts (left-shift) (Figure 3H, I). This was reflected by the fact that *gata2b^-/-^* HSPC1s start expression of *hbba1* (Figure 6G). As the erythroid lineage seems to be stuck in its differentiation, this indicates that Gata2 is important for erythroid lineage differentiation and the initiation of Gata1 expression (Briegel et al., 1993; Moriguchi et al., 2015).

Single cell transcriptome analysis showed further upregulation of genes related to proliferation suggestive of a role for Gata2b in proliferation. Whether this regulation is direct is not yet clear because the lymphoid transcriptional program also regulates cell cycle and this may explain the increase in proliferation signatures in Gata2b deficient HSPCs as they are lymphoid biased.

The role of Gata2 in embryonic generation of HSPCs differs between mammals and the zebrafish system. In mouse and human, Gata2 is required for EHT (de Pater et al., 2013; Huang et al., 2015; Kang et al., 2018), while in zebrafish, *gata2b^-/-^* embryos still show EHT events. High maternal expression of *gata2b* reported earlier by Butko et al., and therefore residual Gata2b protein levels, could possibly rescue EHT. However, maternal zygotic *gata2b* mutants are viable, indicating that embryonic hematopoiesis is normal and maternal expression of *gata2b* does not affect embryonic hematopoiesis (Figure S2B). Previous expression analysis of *gata2a* and *gata2b* indicate that they are both expressed in hemogenic endothelium (Butko et al., 2015; Dobrzycki T, 2019). Therefore, Gata2a could be required for specification of hemogenic endothelium and regulate EHT. The expression of Runx1 is used to mark hemogenic endothelium in the the dorsal aorta (Swiers et al., 2013). In a *gata2a* enhancer mutant (Dobrzycki T, 2019), specifically deleting the expression of *gata2a* in hemogenic endothelium, we found reduced *runx1* expression in *gata2a^i4/i4^* mutant zebrafish (Figure S6A-C). This indicates that endothelial expression of *gata2a* is required upstream of *gata2b* for the proper specification of hemogenic endothelium. In mouse, Gata2 is also required for the maintenance of HSCs after they are generated (de Pater et al., 2013). Why the adult zebrafish compartment is not affected similarly as the mouse HSC compartment by loss of Gata2 is subject for future studies.

In conclusion, we find that Gata2b does not only play an important role in embryonic hematopoiesis, but is vital for the lineage discision between myeloid and lymphoid lineage differentiation. These findings could have great implication for human disease like bone marrow failure syndromes, but also in myelodysplastic syndromes and acute myeloid leukemia caused by mutations in GATA2 (Hahn et al., 2011; Ostergaard et al., 2011; Vinh et al., 2010; Wlodarski et al., 2016).

## Materials and Methods

### Animal husbandry

Zebrafish *(Danio rerio)* were kept under standard conditions. Adult zebrafish were analysed under CCD license AVD 1010020171644. Embryos were collected by natural matings and grown at 28.5C in E3 medium with methylene blue. Heterozygous *Tg(fli1a:eGFP)* (Lawson and Weinstein, 2002), *Tg(−6.0itga2b:EGFP)* (known as CD41:GFP)(Ma et al., 2011), *Tg(mpx:GFP)^i114Tg^* (Ellett et al., 2011), *Tg(mpeg1:GFP)^gl22Tg^* (Renshaw et al., 2006) and *Tg(LCK:eGFP)* (Langenau et al., 2004) embryos were used in this study and all transgenics were analysed as heterozygous.

### Generation of Gata2b mutant zebrafish

Gata2b mutant zebrafish were generated using CRISPR/Cas9 targeting of exon 3. sgRNAs were designed using CHOPCHOP software and prepared according to Gagnon et al. (Gagnon et al., 2014) with minor adjustments. Guide RNAs were generated using the Agilent SureGuide gRNA Synthesis Kit, Cat# 5190-7706. Cas9 protein (IDT) and guide were allowed to form ribonucleoprotein structures (RNPs) at RT and injected in 1 cell stage oocytes. 8 embryos were selected at 24 hpf and lysed for DNA isolation. Heteroduplex mobility PCR analysis was performed to test guide functionality and embryos from the same batch were allowed to grow up. To aid future genotyping we selected mutants by screening F1 for a PCR detectable integration or deletion in exon 3. Sequence verification showed that founder 3 had a 28 nt integration resulting in a frameshift truncating mutation leading to 3 new STOP codons in the third exon. To get rid of additional mutations caused by potential off target effects, founder 3 was crossed to WT for at least 3 generations. All experiments were performed with offspring of founder 3.

### Kidney marrow isolation and analysis

Adult zebrafish were euthanized; WKM removed mechanically using tweezers and dissociated by pipetting in PBS/10% FCS to obtain a single-cell suspension. 7-AAD (7-amino-actinomycin D) 0.5mg/L (BDbiosciences)or DAPI 1mg/L were used for live/dead discrimination. The analysis was performed using FACSAria II (BD)

### Zebrafish embryo cell isolation and sorting

Zebrafish embryos were dechorionated, euthanized and dissociated incubating them in PBS/10% FCS with collagenase I, II and IV each 1:100 (Sigma) for 45 minutes at 37C. During incubation, the embryos were pipetted every 10’ to facilitate the dissociation. FACS analysis and sorting were performed on a FACSAria II (BD) using 7AAD (BD) 0.5mg/L or DAPI (BD) 1mg/L as live/dead marker.

### *In situ hybridization* and analysis

0.003% 1-phenyl-2-thiourea (PTU) treated embryos were fixed O/N with 4% PFA in PBS containing 3% sucrose at appropriate stages and subsequently transferred to MeOH. *ish* has been performed according to Chocron et al.,(Chocron et al., 2007). The *cmyb* probe was a kind gift from Roger Patient. *cmyb* expression was quantified as described previously (Dobrzycki et al., 2018).

### Confocal imaging and analysis

The embryos were anesthetized using tricaine (3-amino benzoic acidethylester) 160mg/L and selected for reporter positivity. The embryos were placed in 0.25% agarose with tricaine 160mg/L. Images were obtained using a Leica SP5 confocal microscope pre-warmed at 28C. The images were acquired using a 4uM stepsize and analyzed using Fiji software (ImageJ)(Schindelin et al., 2012). Quantitation of the CD41:GFP^+^Flt:RFP^+^ cells and EHT events was performed manually, the latter with the aid of ImageJ 3D viewer.

### Opera imaging and analysis

*Tg(mpeg:GFP), Tg(mpx:GFP), Tg(LCK:GFP)* embryos were place in a 96 wells zebrafish arraying plate (ZFplate, Hashimoto Electronic Industry Co. Ltd, Japan) and imaged in tiles covering the complete zebrafish, using a spinning disk confocal high throughput microscope system (Opera Phenix, Perkin Elmer) equipped with a dry 10x objective (NA 0.3). Ambient temperature in the Opera Phenix ranged between 24 - 26°C. GFP was excited using a 488 nm solid state laser and detected at 500 – 550 nm emission in a 20 μm stepsize z-stack. In maximum projection images, GFP positive cells were segmented, counted and analyzed for their GFP intensity and morphology using Harmony software (v 4.8)(Perkin Elmer).

### Bulk RNA sequencing on CD41:GFP^low^ population

RNA was isolated from 50-300 adult zebrafish kidney CD41:GFP^low^ sorted cells in TRIzol (Thermo Fisher Scientific) using the manufacturers protocol and amplified using SMARTer Ultra Low Input RNA kit for sequencing (version 4, Clontech). Sequencing libraries were generated using TruSeq Nano DNA Sample Preparation kits (Illumina), according to the low sample protocol and run on Novaseq 6000 instrument (Illumina). GSEA was performed using the broad institute tool (Mootha et al., 2003; Subramanian et al., 2005).

### Single cell RNA sequencing on whole kidney marrow progenitor population

70.000 single viable cells were sorted from kidney marrows from 2 pooled female *tg(CD41:GFP)* zebrafish. These cells were supplemented with additional CD41:GFP^low^ expressing cells to later detect the HSC compartment. One sample was loaded on the chip and proceeded for single cell barcode labeling by the droplet based 10x genomics machine at a time to reduce cell death. cDNA was prepared using the manufacturers protocol (10x Chromium V2) and sequenced on Novaseq 6000 instrument (Illumina). We used two WT replicates and two *gata2b^-/-^* replicates each sequenced to a depth of 50.000 reads per cell.

### Single cell transcriptome analysis

Seurat V3 was used to analyse the single cell expression data (Butler et al., 2018). The replicates were randomly down-sampled to get an even distribution of WT and *gata2b^-/-^* cell numbers and we performed a quality control excluding doublets (high UMI count and high feature count) or dead cells (mitochondrial reads < 0.08%). For the comparative analysis, first, the replicates of WT and *gata2b^-/-^* animals were individually aligned using achor based integration and then WT and *gata2b^-/-^* cells were aligned to eachother. Clustering was calculated using 15 different dimensions with a resolution of 0.6.

Elbow plot analysis showed that at least 6 dimensions should be used, but marker analysis revealed that the NK population and the T cell population could only be separately distinguished by using 15 dimensions. Clustering was visualized by Uniform Manifold Approximation and Projection (UMAP).

### Statistics

All statistical analysis was carried out in GraphPad Prism 5 (GraphPad Software). Normally distributed data were analyzed using One-way ANOVA with Tukey multiple comparison test. Not normally distributed data were analyzed using Kruskal-Wallis with Dunn test.

## Author contribution

EdP conceived the study; EG, CK, HdL, JP, DB, PvS, MvR, and EB performed experiments; EG, CK, MdJ, RH, KG and EdP analysed results; PF and IT provided resources and EG and EdP wrote the manuscript and IT revised the manuscript.

## Acknowledgements

We thank members of the de Pater, Touw, Raaijmakers and Schneider-Kramann labs for helpful discussions, in particular Drs Raaijmakers and Schneider-Kramann. We thanks Dr Cupedo for careful reading of the manuscript. We thank the Experimental Animal Facility of Erasmus MC for animal husbandry and the Erasmus Optical Imaging Center for confocal microscopy services. This research is supported by the European Hematology Association (junior non clinical research fellowship)(EdP), the Dutch Cancer Foundation KWF/Alpe d’HuZes (SK10321)(EdP) and by the Daniel den Hoed Foundation (IT).

## References

Athanasiadis, E.I., Botthof, J.G., Andres, H., Ferreira, L., Lio, P., and Cvejic, A. (2017). Single-cell RNA-sequencing uncovers transcriptional states and fate decisions in haematopoiesis. Nat Commun 8, 2045.

Bertrand, J.Y., Chi, N.C., Santoso, B., Teng, S., Stainier, D.Y., and Traver, D. (2010a). Haematopoietic stem cells derive directly from aortic endothelium during development. Nature 464, 108–111.

Bertrand, J.Y., Cisson, J.L., Stachura, D.L., and Traver, D. (2010b). Notch signaling distinguishes 2 waves of definitive hematopoiesis in the zebrafish embryo. Blood 115, 2777–2783.

Bertrand, J.Y., Jalil, A., Klaine, M., Jung, S., Cumano, A., and Godin, I. (2005). Three pathways to mature macrophages in the early mouse yolk sac. Blood 106, 3004–3011.

Bertrand, J.Y., Kim, A.D., Teng, S., and Traver, D. (2008). CD41+ cmyb+ precursors colonize the zebrafish pronephros by a novel migration route to initiate adult hematopoiesis. Development 135, 1853–1862.

Boisset, J.C., van Cappellen, W., Andrieu-Soler, C., Galjart, N., Dzierzak, E., and Robin, C. (2010). In vivo imaging of haematopoietic cells emerging from the mouse aortic endothelium. Nature 464, 116–120.

Bottardi, S., Mavoungou, L., Bourgoin, V., Mashtalir, N., Affar el, B., and Milot, E. (2013). Direct protein interactions are responsible for Ikaros-GATA and Ikaros-Cdk9 cooperativeness in hematopoietic cells. Mol Cell Biol 33, 3064–3076.

Briegel, K., Lim, K.C., Plank, C., Beug, H., Engel, J.D., and Zenke, M. (1993). Ectopic expression of a conditional GATA-2/estrogen receptor chimera arrests erythroid differentiation in a hormone-dependent manner. Genes Dev 7, 1097–1109.

Bussmann, J., Bos, F.L., Urasaki, A., Kawakami, K., Duckers, H.J., and Schulte-Merker, S. (2010). Arteries provide essential guidance cues for lymphatic endothelial cells in the zebrafish trunk. Development 137, 2653–2657.

Butko, E., Distel, M., Pouget, C., Weijts, B., Kobayashi, I., Ng, K., Mosimann, C., Poulain, F.E., McPherson, A., Ni, C.W., et al. (2015). Gata2b is a restricted early regulator of hemogenic endothelium in the zebrafish embryo. Development 142, 1050–1061.

Butler, A., Hoffman, P., Smibert, P., Papalexi, E., and Satija, R. (2018). Integrating single-cell transcriptomic data across different conditions, technologies, and species. Nat Biotechnol 36, 411–420.

Chocron, S., Verhoeven, M.C., Rentzsch, F., Hammerschmidt, M., and Bakkers, J. (2007). Zebrafish Bmp4 regulates left-right asymmetry at two distinct developmental time points. Dev Biol 305, 577–588.

Ciau-Uitz, A., Monteiro, R., Kirmizitas, A., and Patient, R. (2014). Developmental hematopoiesis: ontogeny, genetic programming and conservation. Exp Hematol 42, 669–683.

de Pater, E., Kaimakis, P., Vink, C.S., Yokomizo, T., Yamada-Inagawa, T., van der Linden, R., Kartalaei, P.S., Camper, S.A., Speck, N., and Dzierzak, E. (2013). Gata2 is required for HSC generation and survival. J Exp Med 210, 2843–2850.

Dobrzycki, T, K.M., Koyunlar C, Rispoli R, Peulen-Zink J, Gussinklo K, de Pater E, Patient R, Monteiro R (2019). A zebrafish model for MonoMAC syndrome identifies an earlier role for gata2 in haemogenic endothelium programming and generation of haematopoietic stem cells In BioRxiv (BioRxiv).

Dobrzycki, T., Krecsmarik, M., Bonkhofer, F., Patient, R., and Monteiro, R. (2018). An optimised pipeline for parallel image-based quantification of gene expression and genotyping after in situ hybridisation. Biol Open 7.

Driever, W., Solnica-Krezel, L., Schier, A.F., Neuhauss, S.C., Malicki, J., Stemple, D.L., Stainier, D.Y., Zwartkruis, F., Abdelilah, S., Rangini, Z., et al. (1996). A genetic screen for mutations affecting embryogenesis in zebrafish. Development 123, 37–46.

Drissen, R., Buza-Vidas, N., Woll, P., Thongjuea, S., Gambardella, A., Giustacchini, A., Mancini, E., Zriwil, A., Lutteropp, M., Grover, A., et al. (2016). Distinct myeloid progenitor-differentiation pathways identified through single-cell RNA sequencing. Nat Immunol 17, 666–676.

Dykstra, B., Kent, D., Bowie, M., McCaffrey, L., Hamilton, M., Lyons, K., Lee, S.J., Brinkman, R., and Eaves, C. (2007). Long-term propagation of distinct hematopoietic differentiation programs in vivo. Cell Stem Cell 1, 218–229.

Ellett, F., Pase, L., Hayman, J.W., Andrianopoulos, A., and Lieschke, G.J. (2011). mpeg1 promoter transgenes direct macrophage-lineage expression in zebrafish. Blood 117, e49–56.

Gagnon, J.A., Valen, E., Thyme, S.B., Huang, P., Akhmetova, L., Pauli, A., Montague, T.G., Zimmerman, S., Richter, C., and Schier, A.F. (2014). Efficient mutagenesis by Cas9 protein-mediated oligonucleotide insertion and large-scale assessment of single-guide RNAs. PLoS One 9, e98186.

Gao, X., Johnson, K.D., Chang, Y.I., Boyer, M.E., Dewey, C.N., Zhang, J., and Bresnick, E.H. (2013). Gata2 cis-element is required for hematopoietic stem cell generation in the mammalian embryo. J Exp Med 210, 2833–2842.

Gekas, C., and Graf, T. (2013). CD41 expression marks myeloid-biased adult hematopoietic stem cells and increases with age. Blood 121, 4463–4472.

Graf, T., and Enver, T. (2009). Forcing cells to change lineages. Nature 462, 587–594.

Haffter, P., Granato, M., Brand, M., Mullins, M.C., Hammerschmidt, M., Kane, D.A., Odenthal, J., van Eeden, F.J., Jiang, Y.J., Heisenberg, C.P., et al. (1996). The identification of genes with unique and essential functions in the development of the zebrafish, Danio rerio. Development 123, 1–36.

Hahn, C.N., Chong, C.E., Carmichael, C.L., Wilkins, E.J., Brautigan, P.J., Li, X.C., Babic, M., Lin, M., Carmagnac, A., Lee, Y.K., et al. (2011). Heritable GATA2 mutations associated with familial myelodysplastic syndrome and acute myeloid leukemia. Nat Genet 43, 1012–1017.

Herbomel, P., Thisse, B., and Thisse, C. (1999). Ontogeny and behaviour of early macrophages in the zebrafish embryo. Development 126, 3735–3745.

Herrington, F.D., and Nibbs, R.J. (2016). Regulation of the Adaptive Immune Response by the IkappaB Family Protein Bcl-3. Cells 5.

Hoppe, P.S., Schwarzfischer, M., Loeffler, D., Kokkaliaris, K.D., Hilsenbeck, O., Moritz, N., Endele, M., Filipczyk, A., Gambardella, A., Ahmed, N., et al. (2016). Early myeloid lineage choice is not initiated by random PU.1 to GATA1 protein ratios. Nature 535, 299–302.

Huang, K., Du, J., Ma, N., Liu, J., Wu, P., Dong, X., Meng, M., Wang, W., Chen, X., Shi, X., et al. (2015). GATA2(-/-) human ESCs undergo attenuated endothelial to hematopoietic transition and thereafter granulocyte commitment. Cell Regen (Lond) 4, 4.

Jaffredo, T., Gautier, R., Eichmann, A., and Dieterlen-Lievre, F. (1998). Intraaortic hemopoietic cells are derived from endothelial cells during ontogeny. Development 125, 4575–4583.

Johnson, K.D., Kong, G., Gao, X., Chang, Y.I., Hewitt, K.J., Sanalkumar, R., Prathibha, R., Ranheim, E.A., Dewey, C.N., Zhang, J., et al. (2015). Cis-regulatory mechanisms governing stem and progenitor cell transitions. Sci Adv 1, e1500503.

Kang, H., Mesquitta, W.T., Jung, H.S., Moskvin, O.V., Thomson, J.A., and Slukvin, II (2018). GATA2 Is Dispensable for Specification of Hemogenic Endothelium but Promotes Endothelial-to-Hematopoietic Transition. Stem Cell Reports 11, 197–211.

Kauts, M.L., De Leo, B., Rodriguez-Seoane, C., Ronn, R., Glykofrydis, F., Maglitto, A., Kaimakis, P., Basi, M., Taylor, H., Forrester, L., et al. (2018). Rapid Mast Cell Generation from Gata2 Reporter Pluripotent Stem Cells. Stem Cell Reports 11, 1009–1020.

Kim, J., Sif, S., Jones, B., Jackson, A., Koipally, J., Heller, E., Winandy, S., Viel, A., Sawyer, A., Ikeda, T., et al. (1999). Ikaros DNA-binding proteins direct formation of chromatin remodeling complexes in lymphocytes. Immunity 10, 345–355.

Kissa, K., and Herbomel, P. (2010). Blood stem cells emerge from aortic endothelium by a novel type of cell transition. Nature 464, 112–115.

Langenau, D.M., Ferrando, A.A., Traver, D., Kutok, J.L., Hezel, J.P., Kanki, J.P., Zon, L.I., Look, A.T., and Trede, N.S. (2004). In vivo tracking of T cell development, ablation, and engraftment in transgenic zebrafish. Proc Natl Acad Sci U S A 101, 7369–7374.

Laumonnier, Y., Syrovets, T., Burysek, L., and Simmet, T. (2006). Identification of the annexin A2 heterotetramer as a receptor for the plasmin-induced signaling in human peripheral monocytes. Blood 107, 3342–3349.

Lawson, N.D., and Weinstein, B.M. (2002). In vivo imaging of embryonic vascular development using transgenic zebrafish. Dev Biol 248, 307–318.

Ling, K.W., Ottersbach, K., van Hamburg, J.P., Oziemlak, A., Tsai, F.Y., Orkin, S.H., Ploemacher, R., Hendriks, R.W., and Dzierzak, E. (2004). GATA-2 plays two functionally distinct roles during the ontogeny of hematopoietic stem cells. J Exp Med 200, 871–882.

Ma, D., Zhang, J., Lin, H.F., Italiano, J., and Handin, R.I. (2011). The identification and characterization of zebrafish hematopoietic stem cells. Blood 118, 289–297.

Macaulay, I.C., Svensson, V., Labalette, C., Ferreira, L., Hamey, F., Voet, T., Teichmann, S.A., and Cvejic, A. (2016). Single-Cell RNA-Sequencing Reveals a Continuous Spectrum of Differentiation in Hematopoietic Cells. Cell Rep 14, 966–977.

Medvinsky, A., and Dzierzak, E. (1996). Definitive hematopoiesis is autonomously initiated by the AGM region. Cell 86, 897–906.

Mehta, C., Johnson, K.D., Gao, X., Ong, I.M., Katsumura, K.R., McIver, S.C., Ranheim, E.A., and Bresnick, E.H. (2017). Integrating Enhancer Mechanisms to Establish a Hierarchical Blood Development Program. Cell Rep 20, 2966–2979.

Mootha, V.K., Lindgren, C.M., Eriksson, K.F., Subramanian, A., Sihag, S., Lehar, J., Puigserver, P., Carlsson, E., Ridderstrale, M., Laurila, E., et al. (2003). PGC-1alpha-responsive genes involved in oxidative phosphorylation are coordinately downregulated in human diabetes. Nat Genet 34, 267–273.

Moriguchi, T., Suzuki, M., Yu, L., Takai, J., Ohneda, K., and Yamamoto, M. (2015). Progenitor stagespecific activity of a cis-acting double GATA motif for Gata1 gene expression. Mol Cell Biol 35, 805–815.

Muller-Sieburg, C.E., Cho, R.H., Karlsson, L., Huang, J.F., and Sieburg, H.B. (2004). Myeloid-biased hematopoietic stem cells have extensive self-renewal capacity but generate diminished lymphoid progeny with impaired IL-7 responsiveness. Blood 103, 4111–4118.

Nandakumar, S.K., Johnson, K., Throm, S.L., Pestina, T.I., Neale, G., and Persons, D.A. (2015). Low-level GATA2 overexpression promotes myeloid progenitor self-renewal and blocks lymphoid differentiation in mice. Exp Hematol 43, 565–577 e561–510.

North, T.E., Goessling, W., Walkley, C.R., Lengerke, C., Kopani, K.R., Lord, A.M., Weber, G.J., Bowman, T.V., Jang, I.H., Grosser, T., et al. (2007). Prostaglandin E2 regulates vertebrate haematopoietic stem cell homeostasis. Nature 447, 1007–1011.

O’Connell, P.A., Surette, A.P., Liwski, R.S., Svenningsson, P., and Waisman, D.M. (2010). S100A10 regulates plasminogen-dependent macrophage invasion. Blood 116, 1136–1146.

Ostergaard, P., Simpson, M.A., Connell, F.C., Steward, C.G., Brice, G., Woollard, W.J., Dafou, D., Kilo, T., Smithson, S., Lunt, P., et al. (2011). Mutations in GATA2 cause primary lymphedema associated with a predisposition to acute myeloid leukemia (Emberger syndrome). Nat Genet 43, 929–931.

Park, S.M., Cho, H., Thornton, A.M., Barlowe, T.S., Chou, T., Chhangawala, S., Fairchild, L., Taggart, J., Chow, A., Schurer, A., et al. (2019). IKZF2 Drives Leukemia Stem Cell Self-Renewal and Inhibits Myeloid Differentiation. Cell Stem Cell 24, 153–165 e157.

Pinho, S., Marchand, T., Yang, E., Wei, Q., Nerlov, C., and Frenette, P.S. (2018). Lineage-Biased Hematopoietic Stem Cells Are Regulated by Distinct Niches. Dev Cell 44, 634–641 e634.

Renshaw, S.A., Loynes, C.A., Trushell, D.M., Elworthy, S., Ingham, P.W., and Whyte, M.K. (2006). A transgenic zebrafish model of neutrophilic inflammation. Blood 108, 3976–3978.

Rodrigues, N.P., Boyd, A.S., Fugazza, C., May, G.E., Guo, Y., Tipping, A.J., Scadden, D.T., Vyas, P., and Enver, T. (2008). GATA-2 regulates granulocyte-macrophage progenitor cell function. Blood 112, 4862–4873.

Rodrigues, N.P., Janzen, V., Forkert, R., Dombkowski, D.M., Boyd, A.S., Orkin, S.H., Enver, T., Vyas, P., and Scadden, D.T. (2005). Haploinsufficiency of GATA-2 perturbs adult hematopoietic stem-cell homeostasis. Blood 106, 477–484.

Sanjuan-Pla, A., Macaulay, I.C., Jensen, C.T., Woll, P.S., Luis, T.C., Mead, A., Moore, S., Carella, C., Matsuoka, S., Bouriez Jones, T., et al. (2013). Platelet-biased stem cells reside at the apex of the haematopoietic stem-cell hierarchy. Nature 502, 232–236.

Sawai, C.M., Babovic, S., Upadhaya, S., Knapp, D., Lavin, Y., Lau, C.M., Goloborodko, A., Feng, J., Fujisaki, J., Ding, L., et al. (2016). Hematopoietic Stem Cells Are the Major Source of Multilineage Hematopoiesis in Adult Animals. Immunity 45, 597–609.

Schindelin, J., Arganda-Carreras, I., Frise, E., Kaynig, V., Longair, M., Pietzsch, T., Preibisch, S., Rueden, C., Saalfeld, S., Schmid, B., et al. (2012). Fiji: an open-source platform for biological-image analysis. Nat Methods 9, 676–682.

Snow, J.W., Trowbridge, J.J., Fujiwara, T., Emambokus, N.E., Grass, J.A., Orkin, S.H., and Bresnick, E.H. (2010). A single cis element maintains repression of the key developmental regulator Gata2. PLoS Genet 6, e1001103.

Subramanian, A., Tamayo, P., Mootha, V.K., Mukherjee, S., Ebert, B.L., Gillette, M.A., Paulovich, A., Pomeroy, S.L., Golub, T.R., Lander, E.S., et al. (2005). Gene set enrichment analysis: a knowledge-based approach for interpreting genome-wide expression profiles. Proc Natl Acad Sci U S A 102, 15545–15550.

Swiers, G., Baumann, C., O’Rourke, J., Giannoulatou, E., Taylor, S., Joshi, A., Moignard, V., Pina, C., Bee, T., Kokkaliaris, K.D., et al. (2013). Early dynamic fate changes in haemogenic endothelium characterized at the single-cell level. Nat Commun 4, 2924.

Tamplin, O.J., Durand, E.M., Carr, L.A., Childs, S.J., Hagedorn, E.J., Li, P., Yzaguirre, A.D., Speck, N.A., and Zon, L.I. (2015). Hematopoietic stem cell arrival triggers dynamic remodeling of the perivascular niche. Cell 160, 241–252.

Tang, Q., Iyer, S., Lobbardi, R., Moore, J.C., Chen, H., Lareau, C., Hebert, C., Shaw, M.L., Neftel, C., Suva, M.L., et al. (2017). Dissecting hematopoietic and renal cell heterogeneity in adult zebrafish at single-cell resolution using RNA sequencing. J Exp Med 214, 2875–2887.

Traver, D., Paw, B.H., Poss, K.D., Penberthy, W.T., Lin, S., and Zon, L.I. (2003). Transplantation and in vivo imaging of multilineage engraftment in zebrafish bloodless mutants. Nat Immunol 4, 1238–1246.

Tsai, F.Y., Keller, G., Kuo, F.C., Weiss, M., Chen, J., Rosenblatt, M., Alt, F.W., and Orkin, S.H. (1994). An early haematopoietic defect in mice lacking the transcription factor GATA-2. Nature 371, 221–226.

Velten, L., Haas, S.F., Raffel, S., Blaszkiewicz, S., Islam, S., Hennig, B.P., Hirche, C., Lutz, C., Buss, E.C., Nowak, D., et al. (2017). Human haematopoietic stem cell lineage commitment is a continuous process. Nat Cell Biol 19, 271–281.

Vinh, D.C., Patel, S.Y., Uzel, G., Anderson, V.L., Freeman, A.F., Olivier, K.N., Spalding, C., Hughes, S., Pittaluga, S., Raffeld, M., et al. (2010). Autosomal dominant and sporadic monocytopenia with susceptibility to mycobacteria, fungi, papillomaviruses, and myelodysplasia. Blood 115, 1519–1529.

Warga, R.M., Kane, D.A., and Ho, R.K. (2009). Fate mapping embryonic blood in zebrafish: multi- and unipotential lineages are segregated at gastrulation. Dev Cell 16, 744–755.

Wlodarski, M.W., Hirabayashi, S., Pastor, V., Stary, J., Hasle, H., Masetti, R., Dworzak, M., Schmugge, M., van den Heuvel-Eibrink, M., Ussowicz, M., et al. (2016). Prevalence, clinical characteristics, and prognosis of GATA2-related myelodysplastic syndromes in children and adolescents. Blood 127, 1387–1397; quiz 1518.

Yamamoto, R., Morita, Y., Ooehara, J., Hamanaka, S., Onodera, M., Rudolph, K.L., Ema, H., and Nakauchi, H. (2013). Clonal analysis unveils self-renewing lineage-restricted progenitors generated directly from hematopoietic stem cells. Cell 154, 1112–1126.

Zon, L.I., Yamaguchi, Y., Yee, K., Albee, E.A., Kimura, A., Bennett, J.C., Orkin, S.H., and Ackerman, S.J. (1993). Expression of mRNA for the GATA-binding proteins in human eosinophils and basophils: potential role in gene transcription. Blood 81, 3234–3241.

