## Supplementary information for "Single cell transcriptome analysis reveals an essential role for Gata2b in hematopoietic lineage decisions in zebrafish" (from another version of this preprint: version 1, 2019-09-01)

Supplementary information Gioacchino et al.,

STAR Methods

*single cell transcriptome analysis of the progenitor compartment of WT and gata2b$^{-/-}$ zebrafish allows for unbiased lineage investigation*

Kidney marrow from 2 female *Tg(CD41:GFP)* WT or *gata2b$^{-/-}$* zebrafish were pooled and sorted (Figure S4A-C). In total 70.000 cells from the gate in panel B were sorted and between 300 and 1900 CD41:GFP$^{low}$ cells were added to this pool in PBS/10%FSC/2%BSA/2% carp serum. The sorting strategy also included the CD41:GFP$^{high}$ expressing cells (Figure S4C) so for every CD41:GFP$^{low}$ cell, 1.9 CD41:GFP$^{high}$ cells were present in our final population. 7630 cells in WT1, 4033 in WT2, 3675 in *gata2b$^{-/-}$* 1 and 5229 in *gata2b$^{-/-}$* 2 were obtained after quality control, with a read depth of approximately 50.000 reads per cell. Gross differences in cell numbers between WT and *gata2b$^{-/-}$* cells may influence cluster identification when a nearest-neighbor algorithm is used. Therefore, all replicates were randomly down-sampled to 3675 to match each other. First WT and Gata2b deficient cells from the replicates were aligned using anchor based integration and then the WT and *gata2b$^{-/-}$* samples were aligned using the same method to correctly identify the clusters, avoiding batch specific differences. We could identify 20 different cell clusters using the R Seurat package (Butler et al., 2018)(Figure S4D).

*Cluster identification*

Using the FindMarkers function in Seurat, differentially expressed genes were identified compared to the other clusters. Subsequently the functions FeaturePlot and VlnPlot were used to analyse gene expression patterns between clusters to test validity and exclusivity of individual clusters. Final marker analysis was visualized using the DotPlot function.

Erythroid cells, thrombocytes, neutrophils, monocytes and their progenitors were identified by comparing our data to known single cell expression analysis and known lineage markers (Athanasiadis et al., 2017; Carmona et al., 2017; Macaulay et al., 2016; Moore et al., 2016; Tang et al., 2017) (Figure S4E). In short, cells of the erythroid lineage are known to express hemoglobins like *hbba1. itga2b* was used as a marker for thrombocytes in the form of *Tg(CD41:GFP)* and *mpeg1.1* for monocytes and *lysozyme (lyz)* for neutrophils. Furthermore, the differentiated populations were devoid of expression of proliferation genes like *myca* and *pcna* indeed suggesting that these are differentiated cells and not a progenitor compartment. Interestingly, one population expressed markers like *CCAAT enhancer binding protein alpha* (*cebpa)* and *granulin1 (grn1)* (Figure S4H) which are expressed in the monocyte lineage. This population also showed very high expression levels of *s100a10b* (Figure S4E). *in situ* hybridization confirmed that macrophages expressed high levels of *s100a10b* (Figure S4F, G).

The sorting strategy for the kidney progenitor population also contains cells of the lymphoid lineage. A lymphoid progenitor population expressing *rag1* was found. Furthermore, this population also expressed the *rag1* homologue *topoisomerase 2a (top2a)* (Figure S4E), probably as an alternative mechanism in V(D)J recombination (Riou et al., 1993), and high expression of proliferation markers *myca* and *pcna* suggestive of the common lymphoid progenitor (CLP). T cell were marked by expression of *tox, il2rb* and *dusp2* expression, NK cells were marked by *nkl.3, nkl.4* and *ccl33.3* and two distinct populations of B cells were marked by expression of *CD37, CD79a* and *pax5*. Interestingly, the second B cell population showed high levels of the immunoglobulins *ighv1-4* and *ighz* (Danilova et al., 2005; Moore et al., 2016) but did not express the proliferative marker *pcna*, indicating this is a more mature or activated population of B cells (Figure S4E). Analysis of IgM:GFP transgenic zebrafish (Page et al., 2013) showed indeed the presence of several B cell populations (data not shown).

One cluster was marked by high expression of *pigrl3.5* which encodes a polymeric Ig receptor(Kortum et al., 2014). The *pigrl3.5* expressing cells are in close association in the UMAP with the NK population suggesting this is a lymphoid population (Figure S4D, E).

A small cluster with expression of *epithelial cell adhesion molecule (epcam)* was detected (Figure S4E). We hypothesize that this may be niche cells, but since this population is very small and our sorting strategy was not meant to obtain the niche cells, we have not further investigated this population.

*HSPC identification*

The most immature population would be an hematopoietic stem or progenitor cell (HSPC). These cell types are marked by little expression of lineage markers, progenitors express proliferation markers and in zebrafish, *fli1a* and *meis1b* are typical HSC markers (Athanasiadis et al., 2017; Macaulay et al., 2016; Tang et al., 2017). The HSPC1 and HSPC2 populations meet these criteria by expressing low levels of lineage markers (Figure S4E, I, J). These populations showed high expression levels of proliferation markers like *myca, pcna* and *mki67* (Figure S4E) and some cells with high levels of the stem cell markers *fli1a* and *meis1b* (Figure S4I, J). Putative HSC, represented by CD41:GFP$^{low}$ sorted cells, are present in both the HSPC1 and HSPC2 clusters (Figure 5D). As expected, the GFP high expressing cells were thrombocytes. *gata2b* gene expression was enriched in the HSPC1 cluster compared to the other clusters (Figure 5E), indicating that this population would be most affected by the loss of Gata2b. Also, highest expression of *fli1a* and *meis1b* were found in HSPC1(Figure S4I, J) indicating that this cluster contains some HSCs.

**Table S1. Related to Figure 3.**

| | WT | *gata2b*$^{-/-}$ |
|---|---|---|
| **Scatter analysis in freq. of single viable cells** | **N = 26** | **N = 23** |
| Erythrocytes in KM (3 to 12 months) | 25.4 ± 2.8 | 25.9 ± 3.0 |
| Progenitors in KM (3 to 12 months) | 26.8 ± 2.0 | 24.4 ± 1.9 |
| Lymphocytes and HSCs in KM (3 to 12 months ) | 18.1 ± 1.1 | 29.0 ± 1.9**** |
| Myeloid cells in KM (3 to 12 months) | 20.3 ± 1.4 | 10.3 ± 0.9**** |
| **Reporter expression in freq. of single viable cells** | | |
| Tg(*mpx*:GFP) (GFP+ in single live cells) | 17.2 ± 3.0 (n=8) | 5.1 ± 1.0 (n=7)** |
| Tg(*mpeg*:GFP)$^{gl22Tg}$ (GFP+ in single live cells) | 3.5 ± 1.9 (n=4) | 2.1 ± 0.7(n=6) |
| Tg(*LCK*:GFP) (GFP+ in single live cells) | 3.1 ± 0.8 (n =4) | 3.4 ± 0.6 (n=4) |
| Tg(CD41 :GFP) (GFP low in single live cells) | 0.4 ± 0.1 (n=9) | 0.4 ± 0.1 (n=8) |
| **Morphology cell type assessment in freq.** | **N = 6** | **N = 6** |
| blasts | 4.0 ± 1.0 | 1.9 ± 0.6 |
| erythroblasts | 51.4 ± 5.1 | 72.5 ± 4.0* |
| promyelocytes | 7.6 ± 1.2 | 4.6 ± 1.2 |
| myelocytes | 9.0 ± 1.4 | 4.9 ± 1.4 |
| metamyelocytes | 4.4 ± 1.2 | 2.8 ± 0.5 |
| rod shaped neutrophils | 4.6 ± 1.4 | 0.9 ± 0.1 |
| segmented neutrophils | 3.3 ± 1.2 | 0.4 ± 0.2 |
| eosinophils | 2.6 ± 0.9 | 0.4 ± 0.2 |
| monocytes | 2.0 ± 0.4 | 0.8 ± 0.3 |
| lymphocytes | 11.2 ± 1.2 | 10.9 ± 2.7 |

The data are mean± SEM. KM; kidney marrow, n= number of zebrafish used in analysis. *P< 0.05, **P< 0.01, ***P<0.001, ****P<0.0001. If data are normally distributed we used One-way ANOVA with Tukey post-test. If data are not normally distributed we used Kruskal-Wallis with Dunn's post-test. For the morphology quantitation a minimum of 200 erythroblasts was assessed per sample.

### Supplementary figure legends

**Figure S1. Related to Figure 3. Maternal zygotic *gata2b*$^{-/-}$ zebrafish are viable**

A) Distribution of genotypes of matings between *gata2b*$^{+/-}$ and *gata2b*$^{+/-}$ zebrafish. B) Distribution of genotypes of matings between *gata2b*$^{+/-}$ and *gata2b*$^{-/-}$ zebrafish. *gata2b* is highly expressed at the 1-2 cell stage (Butko et al., 2015), indicating maternal contribution of *gata2b* mRNA. During HSC generation

(26-40 hpf), maternal contribution of *gata2b* could provide the embryo with sufficient levels of Gata2b to generate normal numbers of HSPCs. To investigate if maternal contribution of *gata2b* affects the survival of *gata2b*$^{-/-}$ zebrafish, we crossed *gata2b*$^{-/-}$ females to *gata2b*$^{+/-}$ males and vice versa. If *gata2b* maternal expression plays a role in survival of *gata2b*$^{-/-}$ zebrafish, we expect that *gata2b*$^{-/-}$ embryos from *gata2b*$^{-/-}$ females are less likely to survive then embryos derived from *gata2b*$^{-/-}$ males, as these embryos have no maternal contribution of WT *gata2b* mRNA. We found mendelian distribution of both genotypes from both breedings concluding that the maternal contribution of *gata2b* does not affect *gata2b*$^{-/-}$ survival.

**Figure S2. Related to Figure 3. Differentiation markers are not altered in *gata2b*$^{-/-}$ embryos**
A) Representative picture of *Tg(mpeg:GFP)* embryo at 54 hpf B) Quantitation of *mpeg*:GFP$^{+}$ cells in WT, *gata2b*$^{+/-}$ and *gata2b*$^{-/-}$ embryos at 54hpf. C) Representative picture of *Tg(mpx:GFP)* embryos at 75 hpf D) Quantitation of *mpx*:GFP$^{+}$ cells in WT, *gata2b*$^{+/-}$ and *gata2b*$^{-/-}$ embryos. E) Representative picture of *Tg(LCK:GFP)* embryos at 5 dpf F) LCK:GFP$^{+}$ area represented as pixel number in WT, *gata2b*$^{+/-}$ and *gata2b*$^{-/-}$ embryos. Each dot represent one embryo. Error bars represent SEM.

**Figure S3. Related to Figure 4. Transcriptional network analysis indicates decreased expression of genes involved in cellular re-organization in phenotypic HSCs from *gata2b*$^{-/-}$ zebrafish**
A) GSEA analysis indicating enrichment for nuclear chromosome signature, B) cytoskeletal protein binding, C) cell activation, D) chromosomal part, E) establishment of vesicle localization and F) establishment of organelle localization signatures in CD41$^{low}$ cells. Normalized enrichment score, NES.

**Figure S4. Related to Figure 5 and STAR methods. Single cell RNA sequencing reveals several progenitor populations**
A-C) Experimental strategy to obtain single cells for RNA sequencing. B) FACS plot indicating the progenitor population which was sorted and supplemented with the remaining C) CD41:GFP$^{low}$ expressing cells from the kidney marrow pool of cells. D) UMAP showing cluster analysis on aggregated data set of both WT and *gata2b*$^{-/-}$ cells indicating 20 different clusters. E) Dotplot analysis with identifying markers for each of the clusters. F, G) two representative images from *s100a10b in situ* hybridization on zebrafish paraffin sections in the area of the kidney marrow with macrophages indicated with the arrow and enlarged in the corner. Scale bar indicates 10 µm. H) feature analysis of *grn1* expression in blue. Note the expression in the s100a10b clusters. I, J) Violin plots of H) *fli1a* expression and I) *meis1b* expression within the different clusters. Each dot represents the expression level in one cell. UMAP = Uniform manifold approximation and Projection.

**Figure S5. Related to Figure 6. Reclustering of HSPC2 shows high expression of proliferation and lymphoid genes while less expression of myeloid genes**
A) Cluster selection for reclustering HSPC2. B) Reclustering of the HSPC2 population split between WT and *gata2b*$^{-/-}$cells. C) Genotype distribution of WT and *gata2b*$^{-/-}$ HSPC2 in each of the clusters with WT cells in white and *gata2b*$^{-/-}$cells in black. D-Q) Violin plots of marker expression in the different clusters split by genotype with expression in WT cells in green and expression in *gata2b*$^{-/-}$cells in pink. D) proliferation marker *mki67*, E) *myca*, F) *pcna.* Stem cell marker expression G) *fli1a.* Lymphoid marker

expression H) *ikzf1* I) *Fc receptor, IgE, high affinity I, gamma polypeptide like* (*fcer1gl*) J) *ighv1-4* K) *ccr9a* L) *xbp1*. Myeloid lineage marker expression M) *s100a10b* N) *cebpb* O) *cebpa* P) *hbaa2* Q) *ppt1*.

**Figure S6. Related to discussion. *gata2a*$^{i4/i4}$ embryos show decreased *runx1* signal in ISH at 26hpf**
A) Quantitation of signal intensity relative to background cells in WT and *gata2*$^{i4/i4}$ embryos at 26 hpf where each dot represents one embryo B) Representative example of *runx1* expression in WT and *gata2*$^{i4/i4}$ embryos at 26 hpf in the AGM region. *** = P < 0.001, error bars represent SEM.

**Supplementary references**

Athanasiadis, E.I., Botthof, J.G., Andres, H., Ferreira, L., Lio, P., and Cvejic, A. (2017). Single-cell RNA-sequencing uncovers transcriptional states and fate decisions in haematopoiesis. Nat Commun *8*, 2045.
Butko, E., Distel, M., Pouget, C., Weijts, B., Kobayashi, I., Ng, K., Mosimann, C., Poulain, F.E., McPherson, A., Ni, C.W*., et al.* (2015). Gata2b is a restricted early regulator of hemogenic endothelium in the zebrafish embryo. Development *142*, 1050-1061.
Butler, A., Hoffman, P., Smibert, P., Papalexi, E., and Satija, R. (2018). Integrating single-cell transcriptomic data across different conditions, technologies, and species. Nat Biotechnol *36*, 411-420.
Carmona, S.J., Teichmann, S.A., Ferreira, L., Macaulay, I.C., Stubbington, M.J., Cvejic, A., and Gfeller, D. (2017). Single-cell transcriptome analysis of fish immune cells provides insight into the evolution of vertebrate immune cell types. Genome Res *27*, 451-461.
Danilova, N., Bussmann, J., Jekosch, K., and Steiner, L.A. (2005). The immunoglobulin heavy-chain locus in zebrafish: identification and expression of a previously unknown isotype, immunoglobulin Z. Nat Immunol *6*, 295-302.
Kortum, A.N., Rodriguez-Nunez, I., Yang, J., Shim, J., Runft, D., O'Driscoll, M.L., Haire, R.N., Cannon, J.P., Turner, P.M., Litman, R.T*., et al.* (2014). Differential expression and ligand binding indicate alternative functions for zebrafish polymeric immunoglobulin receptor (pIgR) and a family of pIgR-like (PIGRL) proteins. Immunogenetics *66*, 267-279.
Macaulay, I.C., Svensson, V., Labalette, C., Ferreira, L., Hamey, F., Voet, T., Teichmann, S.A., and Cvejic, A. (2016). Single-Cell RNA-Sequencing Reveals a Continuous Spectrum of Differentiation in Hematopoietic Cells. Cell Rep *14*, 966-977.
Moore, J.C., Tang, Q., Yordan, N.T., Moore, F.E., Garcia, E.G., Lobbardi, R., Ramakrishnan, A., Marvin, D.L., Anselmo, A., Sadreyev, R.I*., et al.* (2016). Single-cell imaging of normal and malignant cell engraftment into optically clear prkdc-null SCID zebrafish. J Exp Med *213*, 2575-2589.
Page, D.M., Wittamer, V., Bertrand, J.Y., Lewis, K.L., Pratt, D.N., Delgado, N., Schale, S.E., McGue, C., Jacobsen, B.H., Doty, A*., et al.* (2013). An evolutionarily conserved program of B-cell development and activation in zebrafish. Blood *122*, e1-11.
Riou, J.F., Grondard, L., Petitgenet, O., Abitbol, M., and Lavelle, F. (1993). Altered topoisomerase I activity and recombination activating gene expression in a human leukemia cell line resistant to doxorubicin. Biochem Pharmacol *46*, 851-861.
Tang, Q., Iyer, S., Lobbardi, R., Moore, J.C., Chen, H., Lareau, C., Hebert, C., Shaw, M.L., Neftel, C., Suva, M.L*., et al.* (2017). Dissecting hematopoietic and renal cell heterogeneity in adult zebrafish at single-cell resolution using RNA sequencing. J Exp Med *214*, 2875-2887.

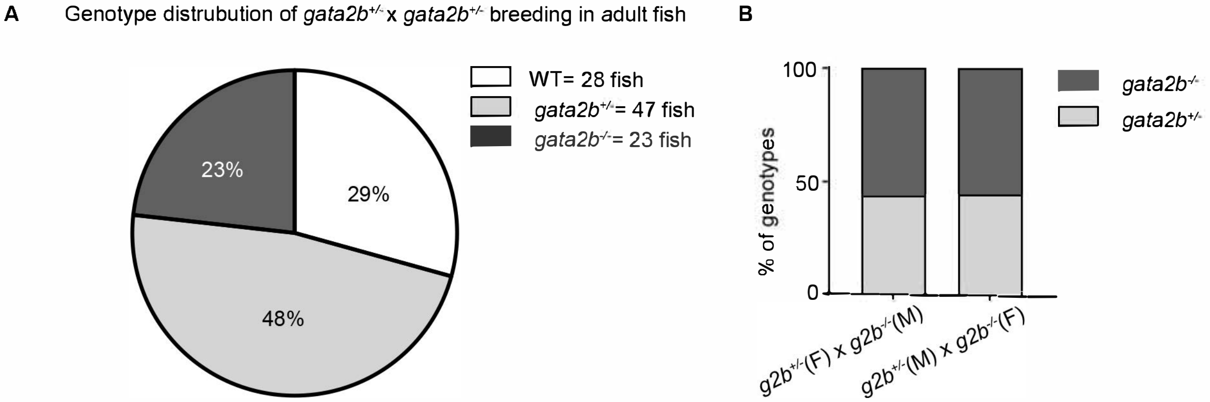


**Figure S1. Related to Figure 3**

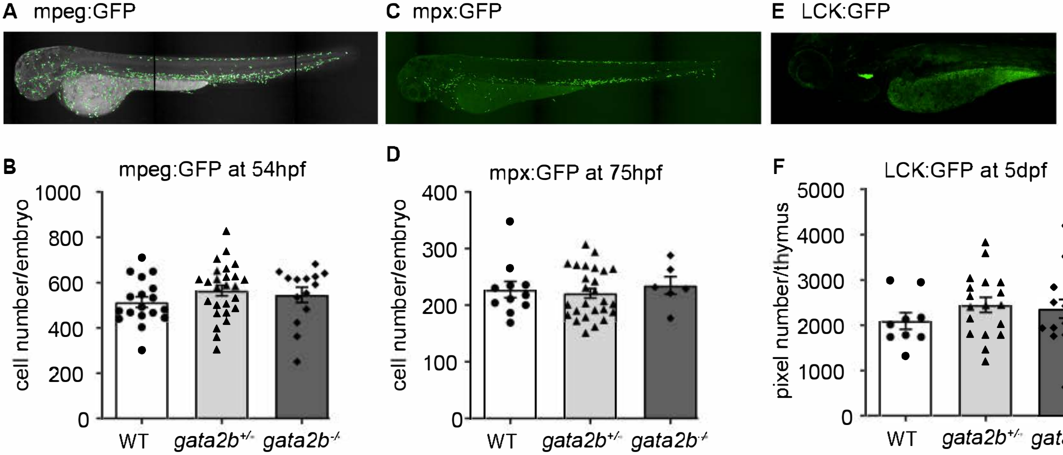


**Figure S2. Related to Figure 3**

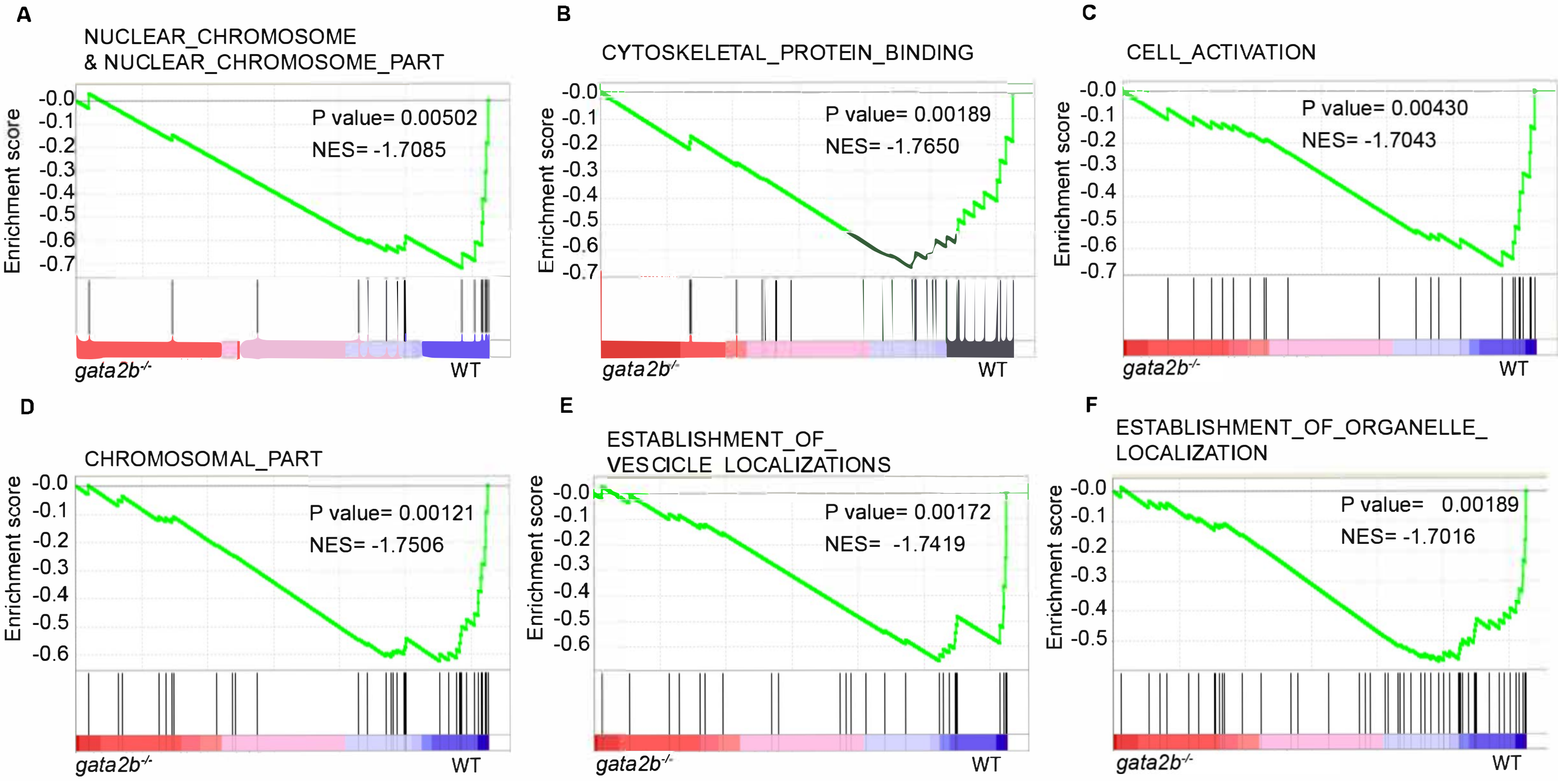


**Figure S3. Related to Figure 4**

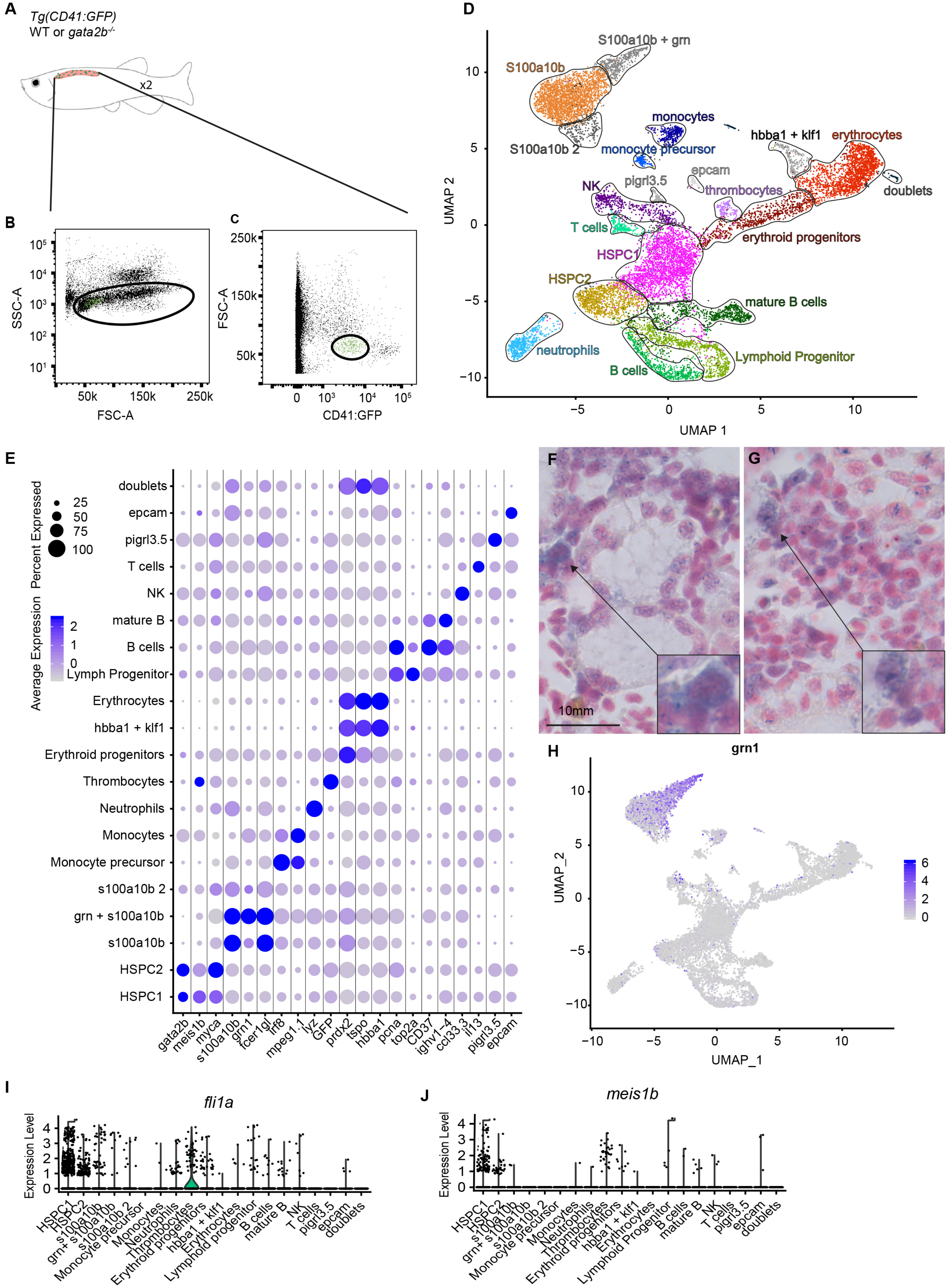


**Figure S4. Related to Figure 5.**

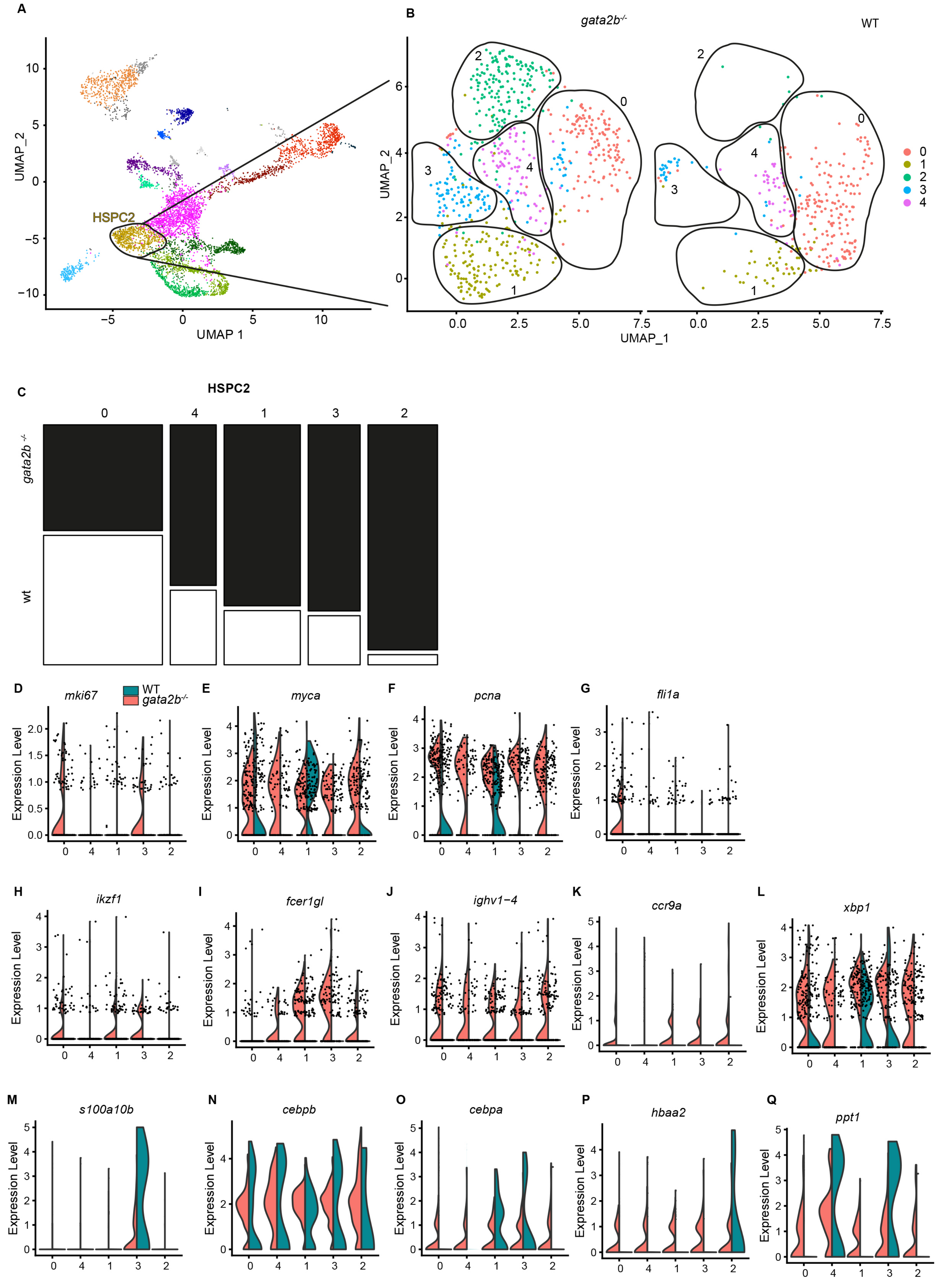


**Figure S5. Related to Figure 6 .**

**A**

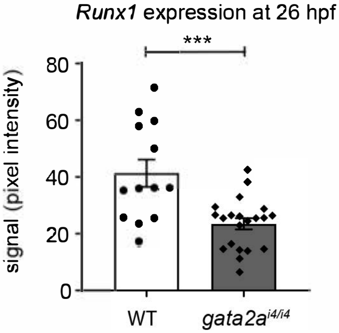


**B**

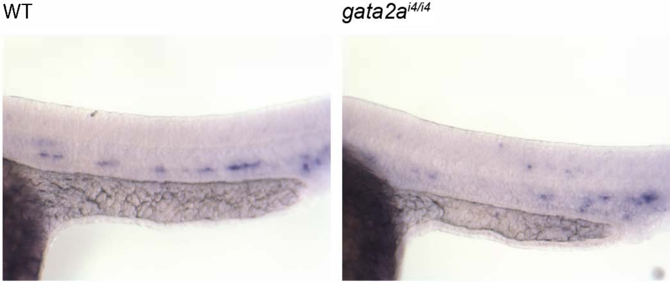


**Figure S6. Related to discussion**
